# Linguistic Context Shapes Representations in the Human Visual Cortex

**DOI:** 10.1101/2025.11.03.686222

**Authors:** Shurui Li, Zheyu Jin, Ru-Yuan Zhang, Shi Gu, Yuanning Li

## Abstract

Language provides a way to anticipate visual scenes before they are seen, but how this context relates to visual representations across levels of abstraction remains unclear. Here, 210 hours of functional magnetic resonance imaging from eight participants performing a delayed caption-image matching task are analyzed using computational encoding, decoding, and alignment models. Caption-evoked activity carries semantic information across visual cortex, including early visual areas. Subsequent image responses vary with both the strength of caption-derived contextual representations and the discrepancy between anticipated and viewed scenes, assessed across lower-level visual and higher-level semantic features. Two functionally distinct voxel populations are identified within early visual cortex: suppression populations respond less strongly to matching than mismatching images, whereas enhancement populations show the opposite pattern. Despite their lower responses to matching images, suppression populations support comparable or better retrieval of corresponding caption-evoked responses using fewer voxels. The two populations also differ in their temporal dynamics, representational structure, and relationship to behavior. Together, these findings link linguistic context to both the strength and informational content of visual cortical responses, revealing how language-vision relationships are expressed across representational levels and two functionally distinct populations in early visual cortex.

## Introduction

Perception is not determined by sensory input alone^1–3^. Visual processing is continuously shaped by contextual information that helps specify what is relevant, likely, or meaningful in a given moment^4–6^. Among the many forms of context that shape perception, language is especially powerful because it can establish rich scene-level meaning before visual input arrives^7,8^. A brief verbal description can specify likely objects, relations, and the overall gist of an upcoming scene^9–11^, thereby linking abstract semantic knowledge to later sensory processing. Understanding how such language-defined context influences visual processing is therefore central to explaining how knowledge and perception interact during natural scene understanding.

Previous work has shown that linguistic context and other top-down factors can modulate activity in visual cortex^12–21^. However, the key unresolved issue is not simply whether language can influence visual processing, but how abstract, language-defined semantic context is transformed into changes in visual representations during natural scene perception^22–24^. Language differs from simple cues or visually defined expectations in an important way: it conveys relational and scene-level meaning without specifying a precise sensory template. As a result, any preparatory activity elicited before image onset need not reflect a single mechanism such as explicit mental imagery, but may instead arise from a broader family of language-driven contextual processes that organize or bias subsequent visual processing. It therefore remains unclear whether language-defined context primarily affects later interpretive stages, or whether it is also expressed within visual representations themselves, including in early visual cortex^19,25^.

This question is especially important for natural scene perception, where language can constrain the semantic structure of an upcoming scene across multiple representational levels^26–28^. If language provides structured semantic constraints rather than a simple global bias, its influence on visual cortex may likewise be structured rather than uniform^29–33^. In that case, contextual effects may be implemented through distinct neural populations that differ in how they relate prior semantic context to incoming visual input. A central question, therefore, is whether language-defined scene meaning is reflected in cortical activity before image onset, whether the strength and structure of caption-image association shape subsequent visual responses, and whether these effects are organized through heterogeneous neural populations rather than distributed uniformly across cortex.

Here we addressed these questions using a large-scale fMRI dataset collected during a delayed caption-image matching task. By presenting a natural-language caption^34^ before a natural scene image^35^, this design allowed us to dissociate caption-evoked activity from image-evoked activity and to examine how visual processing unfolded under an already established language-defined context. We combined model-based analyses of semantic information and graded caption-image relationships with population-level analyses to test whether language-defined context is expressed within visual cortex itself, whether it shapes later responses to visual input, and whether these effects are organized at the level of distinct neural populations.

We found that language-defined context shapes visual processing through linked effects across cortical scales. Before image onset, caption-evoked activity carried structured semantic information across visual cortex, including early visual cortex, indicating that language can establish contextual states related to upcoming scene meaning. After image onset, early visual responses were jointly explained by the strength of these contextual states and model-derived caption-image semantic discrepancy. Thus, visual activity reflected not only incoming sensory input, but also the semantic structure imposed by prior linguistic context. This discrepancy-related structure was distributed across early visual areas and weakened when contextual statistics were altered, indicating that it depended on the predictive structure of the task. Within early visual cortex, language-driven contextual effects were not uniform. Suppression- and enhancement-defined populations differed in cross-modal alignment, temporal dynamics, covariance structure, behavioral coupling and semantic readout. Together, these findings show that language-defined scene context is embedded in the functional organization of visual cortex, shaping visual representations through context-dependent interactions among distinct neural populations rather than through a single global top-down bias.

## Results

### Overview

In all experiments, subjects first read a caption and then viewed a natural image from COCO-CN^34^, judging whether the caption and image matched (Fig. 1a). This design allowed us to dissociate caption-evoked from image-evoked responses, such that activity during caption presentation could be examined before visual input, whereas activity during image viewing could be interpreted in relation to the semantic context established by the preceding caption. The separability of these two response components was further supported by supplementary analyses. Specifically, Supplementary Fig. 1 shows GLM fits from two example voxels, illustrating dissociable caption- and image-evoked response profiles, whereas Supplementary Fig. 2-4 provide additional evidences that adjacent caption- and image-related signals were separable at the population level. To test whether different contextual regimes shape subsequent visual responses, we further conducted a control experiment (Fig. 1b) in which the prevalence of matched and mismatched trials was reversed relative to the main experiment. This manipulation altered the overall contextual statistics while preserving the same task format, allowing us to compare image-evoked responses under different expectancy structures. In addition, eye-movement heatmaps indicated stable viewing behavior and sustained task engagement throughout the experiment (Supplementary Fig. 5). Across eight subjects, we collected approximately 210 hours of 3T fMRI data, enabling model-based analyses of how language-defined scene context relates to responses in visual cortex.

**Figure 1.**
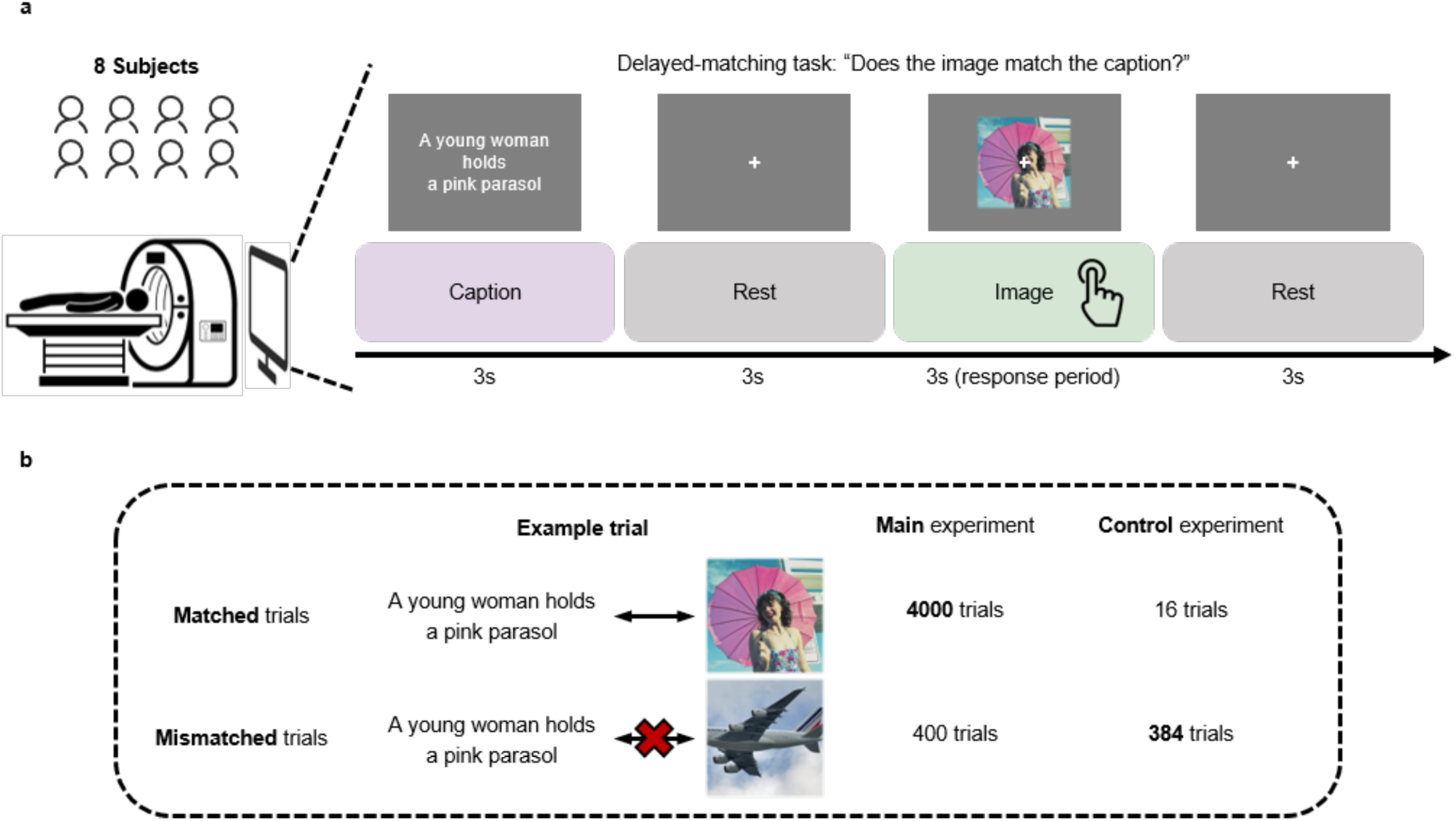
Experimental paradigm. **a)** Temporally segregated delayed-matching paradigm. Each trial lasted 12 seconds: participants first viewed a caption for 3 s, followed by a 3 s blank interval, a 3 s natural image, and a final 3 s interval. During the image period, participants judged whether the image matched the preceding caption, and responded only when they detected a mismatch. **b)** Trial composition and manipulation of prior information. In the main experiment, 4,000 matched and 400 mismatched trials were collected per subject (10:1 ratio), whereas in the control experiment, 32 matched and 384 mismatched trials were collected (2:23 ratio) to assess the influence of prior expectations on visual responses. See Methods for full details.

### Semantic priors reshape responses in early visual cortex

We first asked whether image-evoked responses in early visual cortex depended on the semantic relationship between a preceding caption and the viewed image. In the main experiment, captions established scene-level expectations several seconds before image presentation, allowing us to contrast responses to semantically matched and mismatched caption-image pairs.

We analyzed image-evoked responses from three complementary perspectives. First, using single-trial response estimates from an event-related GLM, we performed voxel-wise contrasts between matched and mismatched conditions. We found that responses during matched trials were consistently lower than those during mismatched trials, indicating significant suppression. Cluster-based statistical testing (p < 0.05, cluster size > 30) confirmed that this suppression was robust across all eight participants, predominantly in EVC and along the ventral visual stream (Fig. 2a). We further visualized the population receptive fields (pRFs) of suppression voxels and found that they were concentrated within the visual field region occupied by the image, whereas other visually responsive voxels showed a less spatially confined distribution (Supplementary Fig. 6). To assess group-level consistency, we projected individual maps onto an averaged cortical surface and computed the proportion of participants exhibiting significant matched–mismatched differences at each vertex. Using a subject index threshold of 0.25, we observed a spatially coherent pattern spanning EVC and extending anteriorly along ventral occipitotemporal cortex, indicating that caption-defined semantic context is associated with reliable modulation of visual responses in early stages of visual processing. Although some enhancement effects were also observed in the dorsal stream (Supplementary Fig. 7), the dominant and most spatially consistent effect in sensory cortex was lower responses for matched than mismatched trials.

**Figure 2.**
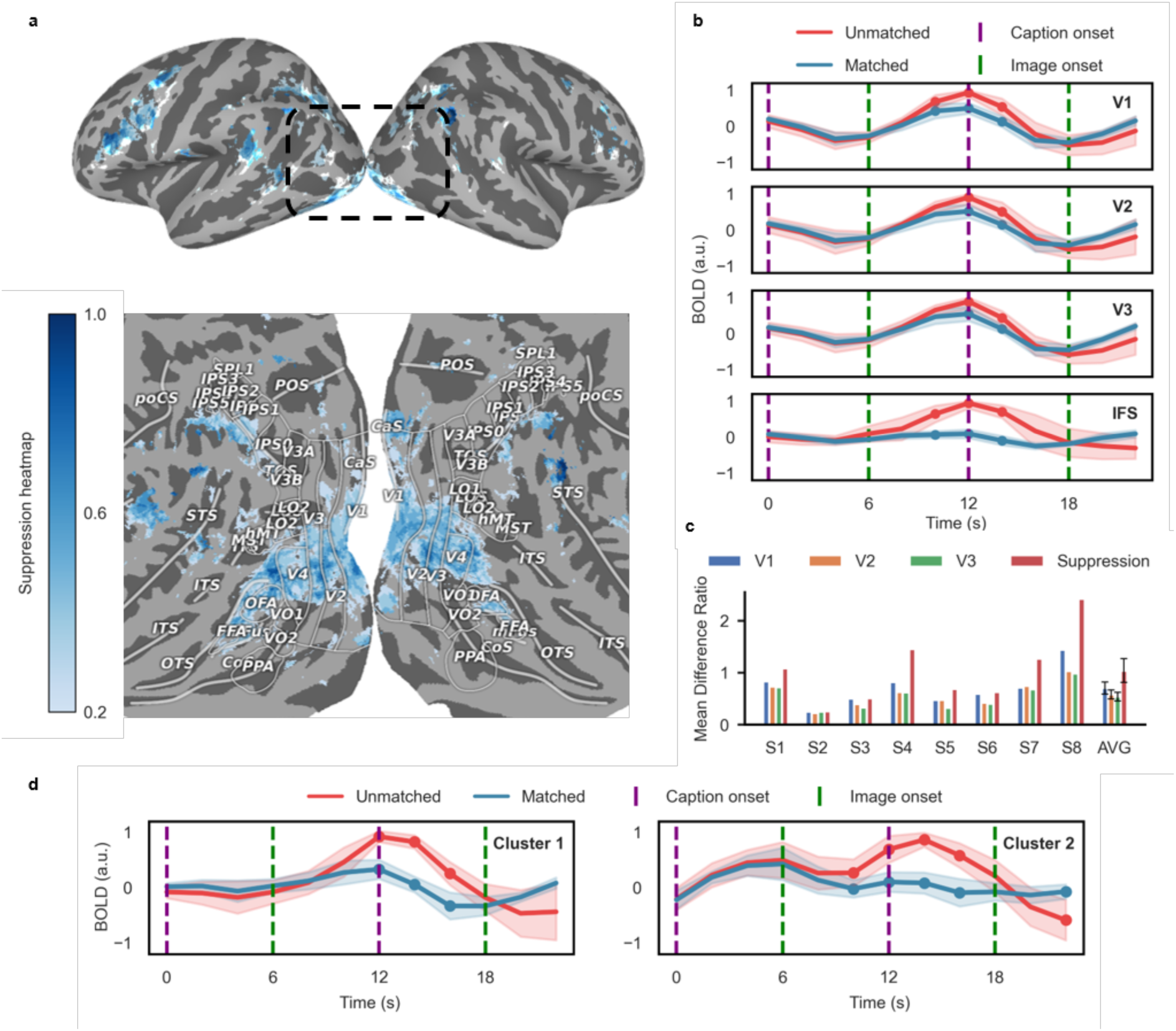
Image-evoked response differences associated with caption-defined semantic context. **a)** Group-level distribution of suppression voxels across the whole brain. The boxed region highlights the ventral visual cortex, with a magnified view shown below. The suppression heatmap reflects the number of participants exhibiting suppression at each voxel after alignment to the standard cortical template. **b)** Averaged normalized time courses (mean ± s.d., across 8 participants) within four ROIs (V1, V2, V3, and IFS), plotted for matched and unmatched trials. Time series are scaled to [−1, 1]. Dots indicate time points with significant differences between conditions (p < 0.05, Wilcoxon signed-rank test, FDR corrected). **c)** Relative difference in response variance between unmatched and matched trials across 4 ROIs (V1, V2, V3, IFS). “AVG” denotes the group average across 8 participants; error bars indicate standard deviation. **d)** Two representative temporal clusters identified via time series clustering. Cluster 1 shows a canonical visual response to image stimuli, while Cluster 2 exhibits responses to both image and caption stimuli. Dots mark time points with significant differences between matched and unmatched conditions (p < 0.05, Wilcoxon signed-rank test, FDR corrected).

Second, we examined trial-aligned time courses within anatomically defined ROIs in V1, V2, and V3. Across all three ROIs, we observed markedly reduced response amplitudes during match trials of images compared to those in mismatched trials (Fig. 2b). Despite variability in spatial extent across individuals, suppression was robustly present in all participants, further confirming the effect beyond beta estimates. Notably, we also observed a similar suppression pattern in IFS, a higher-order association area involved in top-down control and semantic processing. Moreover, a similar pattern is observed in fusiform face area (FFA), and parahippocampal place area (PPA) (Supplementary Fig. 8). This suggests that semantic prior modulation in multimodal integration extends beyond the visual cortex and reflects a more widespread network-level mechanism supporting multimodal semantic integration.

Thirdly, to assess the overall response variability under different predictive conditions, we computed the standard deviation of the trial-averaged time series within each ROI (Fig. 2c). Across all four regions (V1, V2, V3, and IFS), time-series variability was consistently higher in mismatched than matched trials. Within early visual cortex, the mismatched–matched difference decreased from V1 to V3. This pattern suggests that lower-level visual areas exhibit greater response fluctuations under semantic mismatches, potentially reflecting higher sensitivity to expectation violations. Alternatively, this effect may arise because low-level visual features are inherently more difficult to predict from abstract linguistic cues, resulting in greater residual variability when top-down expectations fail to constrain sensory input. Moreover, the suppression region exhibited significantly higher difference than any of the anatomical ROIs, suggesting that this functionally localized area is particularly sensitive to top-down modulation and reflects enhanced dynamic range under violated expectations.

Finally, to further probe the diversity of neural responses within the suppression regions, we performed voxel-wise clustering based on response time courses. Two dominant voxel types emerged: (1) unimodal voxels with a single peak aligned to image onset, and (2) bimodal voxels responsive to both caption and image events (Fig. 2d). Both voxel types showed lower responses in matched than mismatched trials, indicating that the matched–mismatched difference generalized across distinct response profiles within suppression-defined cortex.

Together, these results show that image-evoked responses in early visual cortex depend systematically on the semantic relationship between a preceding caption and the viewed image. Rather than being uniformly expressed, these effects are detectable across both anatomical ROIs and functionally defined subregions, providing an initial description of language-guided semantic context effects on visual cortical responses.

### Early visual cortex encodes language-derived semantic information

We next asked whether caption-evoked activity contained structured language-derived information before image onset. We tested this using a caption encoding analysis, in which semantic features extracted from pretrained language models were used to predict fMRI responses during caption presentation.

We employed a multimodal encoding framework using CLIP-BERT, a transformer-based language-vision model^36,37^, and extracted contextualized text embeddings from its language encoder to capture rich semantic representations of the captions for neural response prediction. We extracted language features from each text caption across all eight transformer layers, and used a weighted combination of these features to predict voxel-wise fMRI responses to the text caption via ridge regression^38,39^ (Fig. 3a). Model performance was assessed across the entire cortex. As shown in Fig. 3b, the encoding model revealed robust prediction accuracy in classical semantic and category-selective areas, including lateral occipital complex (LOC), inferotemporal cortex (IT), fusiform face area (FFA), and parahippocampal place area (PPA), consistent with previous studies^40–43^. Notably, despite weaker overall performance in EVC, significant prediction was still observed, suggesting the presence of structured language-derived signals in the early visual cortex.

**Figure 3.**
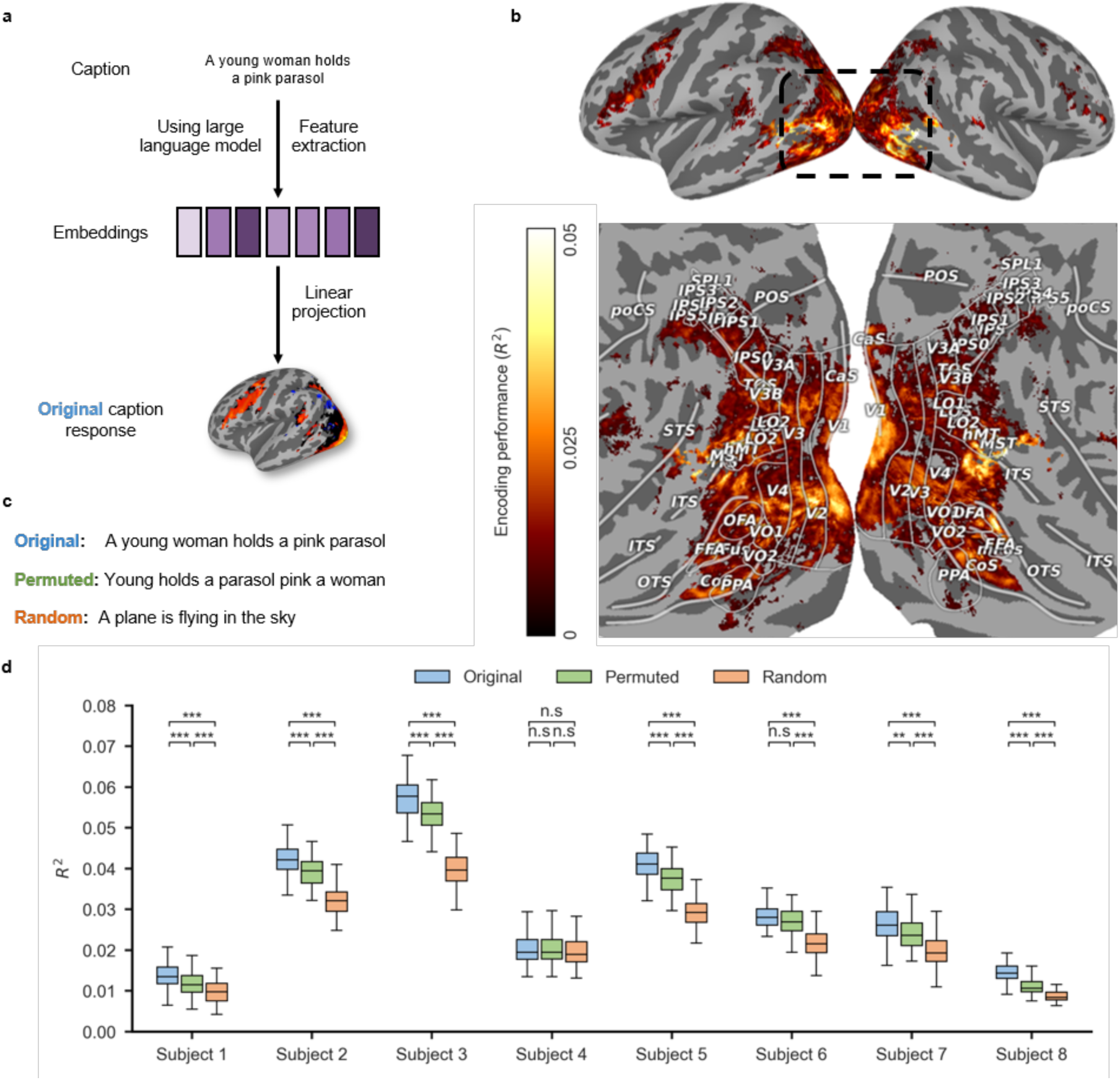
Caption-evoked responses in early visual cortex contain language-derived semantic information. **a)** Schematic of the linear encoding model. Embeddings were extracted from the original caption using a pretrained large language model, followed by feature mixing (not shown). The mixed features were then used to predict voxel-wise brain responses. **b)** Whole-brain performance of the caption encoding model, quantified using *R*^2^. Only voxels with ONOFF-*R*^2^ > 1% are shown. The boxed region highlights early visual cortex, with a magnified view presented below. No noise ceiling normalization was applied. **c)** Examples of the three caption conditions: original refers to the actual text presented to participants; permuted captions were created by shuffling the word order of the original; random captions were replaced with unrelated sentences. **d)** Prediction performance (*R*^2^) across the three caption conditions for each of the 8 participants using CLIP-Bert. Asterisks indicate statistical significance across conditions (n.s.: not significant, *: p < 0.05, **: p < 0.01, ***: p < 0.001; Mann–Whitney U test, Bonferroni correction).

To directly assess the contribution of the semantic content in the text caption, we introduced two control manipulations. In the permuted condition, the order of words within the original caption was shuffled, disrupting sentence-level semantic coherence while preserving lexical and orthographic features but disrupting semantics. In the random condition, the original caption was replaced by a random and syntactically unrelated sentence, removing stimulus-specific semantic content altogether (Fig. 3c). Then we fed the texts of these two conditions to the CLIP-Bert model and used the extracted features to predict the original fMRI caption response. These manipulations allowed us to dissociate contributions from semantic, lexical, and low-level visual properties of the text.

Across all eight participants, we found that model performance across the whole brain was highest for the original captions (averaged *R*^2^ = 0.03044), followed by the permuted condition (averaged *R*^2^ = 0.02808), with the random condition yielding the lowest prediction accuracy (averaged *R*^2^ = 0.02262) (Fig. 3d). Critically, EVC showed the same ordering, with significant reductions in prediction performance when semantic structure was disrupted. Consistently, when RoBERTa^44,45^ was used as the backbone for the encoding model, we obtained a highly similar pattern: averaged *R*^2^ = 0.03214 for the original captions, *R*^2^ = 0.02702 for the permuted condition, and *R*^2^ = 0.02110 for the random condition (Supplementary Fig. 9). This suggests that the full semantic content of the caption, rather than lexical form or orthography alone, contributes to caption-evoked responses, including in early visual cortex.

In sum, these results indicate that language-derived semantic information is detectable in caption-evoked activity before image onset, not only in higher-order association regions but also in early visual cortex. This finding provides evidence that caption-defined scene meaning is reflected in pre-image neural responses in EVC.

### Contextual states and semantic discrepancy jointly explain early visual responses

The preceding analysis showed that caption-evoked activity contained structured semantic information across visual cortex before image onset. We next asked whether these pre-image semantic states were related to the visual response elicited by the subsequent image. Rather than moving directly to voxel-wise analyses, we first tested this relationship at the ROI level in early visual cortex (EVC), where image-evoked responses were robust and where contextual modulation would provide direct evidence that language-derived information influenced early stages of visual processing.

We modelled two complementary quantities for each caption-image pair. The first was prior strength, which indexed how strongly the caption established an expected scene representation before image onset (Fig. 4a). Higher prior strength therefore reflected a more clearly constrained prediction about the upcoming image. The second was high-level prediction-error (PE) strength, which indexed the semantic discrepancy between the caption-defined expectation and the image that was actually presented (Fig. 4c). Higher PE strength therefore reflected a larger mismatch at the level of scene meaning.

**Figure 4.**
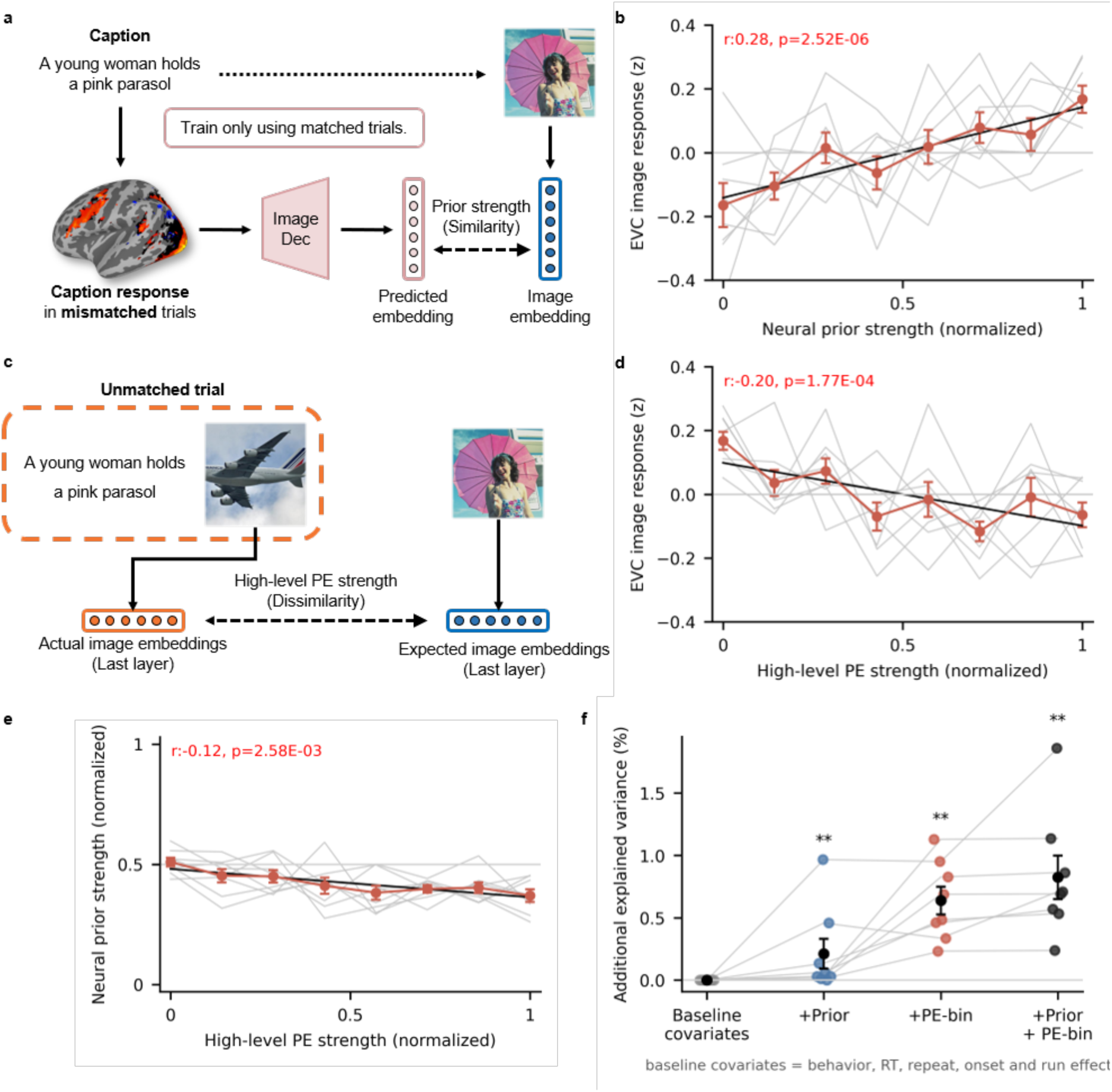
Language-derived priors and semantic discrepancy jointly explain early visual responses. **a)** Schematic of neural prior strength estimation. An image decoder was trained on matched trials to map caption-evoked fMRI responses onto CLIP image embeddings. For mismatched trials, the decoder generated a predicted image embedding from the caption-period response. Neural prior strength was defined as the similarity between this predicted embedding and the CLIP embedding of the caption-consistent image, with higher similarity indicating a more specific language-derived expectation. **b)** Relationship between neural prior strength and image-evoked responses in early visual cortex (EVC). Neural prior strength was normalized to the range 0-1, and EVC responses were z-scored. Trials were divided into eight bins. Grey lines show individual participants, and the red line and shading show the group mean and std. EVC responses increased with neural prior strength (Pearson r = 0.28, p = 2.52e-06). **c)** Schematic of high-level prediction-error (PE) strength estimation. For mismatched trials, high-level PE strength quantified the semantic dissimilarity between the image expected from the caption and the image actually presented. Dissimilarity was computed from final-layer CLIP embeddings, with higher values indicating greater semantic mismatch. **d)** Relationship between high-level PE strength and image-evoked EVC responses. PE strength was normalized to the range 0-1, and EVC responses were z-scored. As in Fig. 4b, trials were divided into eight bins, grey lines show individual participants, and the red line and shading show the group mean and std. EVC responses decreased as high-level PE strength increased (Pearson r = −0.20, p = 1.77e-04). **e)** Relationship between neural prior strength and high-level PE strength across mismatched trials. The two measures were only weakly correlated (Pearson r = −0.12, p = 2.58e-03), indicating that prior strength and semantic discrepancy captured separable components of the caption-image relationship. **f)** Additional variance explained in image-evoked EVC responses after adding neural prior strength, high-level PE strength, or both variables to a baseline model. The baseline model included behavioral response, reaction time, image repetition and presentation time point. Models including prior strength, high-level PE strength, or both variables explained additional variance beyond the baseline model, with the combined model showing the largest improvement (FDR q < 0.01 for all comparisons).

At the ROI level, EVC responses showed reliable but opposite relationships with these two quantities. Image-evoked EVC activity increased with prior strength (Pearson r = 0.28, p = 2.52e-06), indicating that stronger pre-image contextual states were followed by larger responses during image viewing (Fig. 4b). By contrast, EVC activity decreased with high-level PE strength (Pearson r = −0.20, p = 1.77e-04), indicating that larger semantic discrepancy between the expected and viewed image was associated with weaker image-evoked activity (Fig. 4d). Thus, early visual responses were not explained by sensory input alone, but varied systematically with both the strength of the preceding semantic context and its alignment with the incoming image.

We then tested whether prior strength and high-level PE strength captured separable components of the caption-image relationship. The two variables were only weakly correlated (Pearson r = 0.12, p = 2.58e-03), indicating that the clarity of a caption-derived prediction and the semantic discrepancy elicited by the viewed image were related but nonredundant (Fig. 4e). To further assess their contribution, we constructed a baseline model that included behavioural response, reaction time, image repetition and presentation time point. Adding prior strength and high-level PE strength improved the model beyond these baseline covariates (FDR q < 0.01), showing that both variables contributed additional explanatory power for ROI-level EVC responses (Fig. 4f).

Together, these ROI-level analyses indicate that early visual activity reflected two linked aspects of language-defined context: the strength of the pre-image semantic state and the semantic discrepancy between that state and the viewed image. This joint effect provided an initial ROI-level basis for the voxel-wise analyses below, where we asked how contextual state and semantic discrepancy were organized across local response patterns within early visual cortex.

### Image-evoked responses in EVC covary with model-derived semantic discrepancy

We next tested whether image-evoked responses in early visual cortex varied with the magnitude of model-derived semantic discrepancy, beyond categorical caption-image congruency. For mismatched trials, we quantified the feature dissimilarity between the scene implied by the caption and the image that was actually shown, and related these dissimilarity measures to voxel-wise BOLD responses. We restricted this analysis to mismatched trials because they provided a closely balanced comparison between the main and control experiments, which contained similar numbers of trials (400 and 384 respectively). This design minimized differences in statistical power when comparing how contextual structure shaped discrepancy-related responses across protocols.

To capture different levels of visual abstraction, we extracted feature dissimilarity from a range of pretrained convolutional and transformer-based models, including AlexNet^46^, ResNet 18^47^, ResNet 50^47^, and ViT^48^, trained on ImageNet^49^ for general-purpose visual tasks. For each model, we categorized features into three levels: low-level (e.g., early convolutional layers capturing edge and texture information), mid-level (intermediate layers), and high-level (final layers capturing semantic content) feature dissimilarity^50^. Correlation analyses of these feature spaces showed relatively high within-level similarity and lower between-level similarity, supporting this hierarchical grouping (Fig. 5b). In addition, we used CLIP-based text-image dissimilarity as a high-level semantic measure.

**Figure 5.**
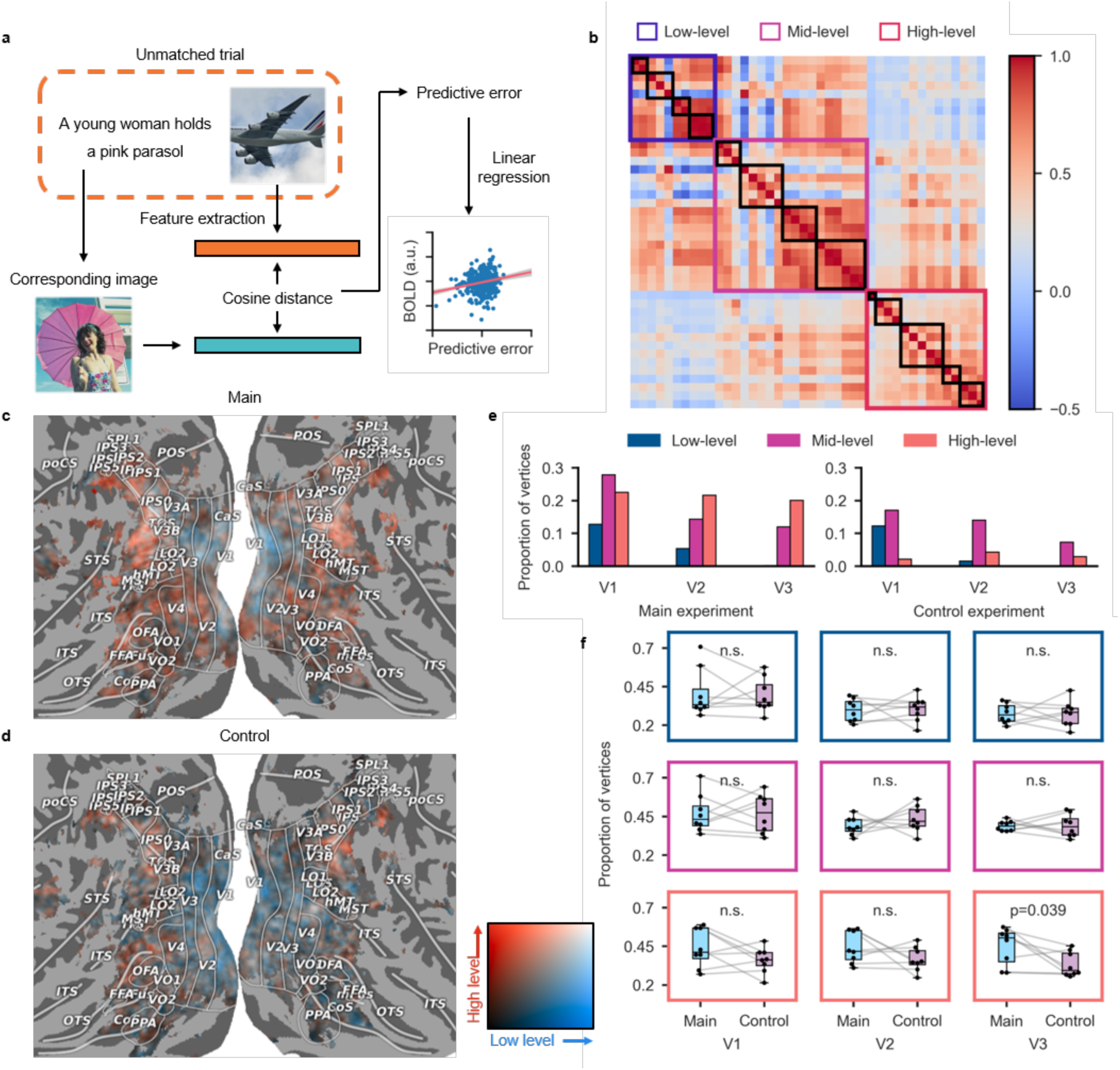
Discrepancy-related feature representations in early visual cortex. **a)** Schematic of the semantic discrepancy model. Visual features were extracted from pretrained vision models. The scatter plot and regression line illustrate an example voxel-wise linear fit (left hemisphere, voxel 449) using features from CLIP-ViT layer 32 (r = 0.21, p = 9.1×10^−5^). **b)** Correlation matrix of image features across all models and layers. Features were empirically grouped into low-, mid-, and high-level representations. Within low- and mid-levels, four subgroups (boxed in black) correspond to CLIP-ViT, AlexNet, ResNet18, and ResNet50. In the high-level group, five subgroups (boxed) correspond to CLIP (image-caption cosine similarity), CLIP-ViT, AlexNet, ResNet18, and ResNet50. **c)** Group-level cortical projection of model-derived semantic discrepancy feature levels in the main experiment. Red indicates voxels correlated with high-level features across multiple participants, blue indicates voxels correlated with low-level features, white indicates voxels correlated with both, and black indicates no significant correlation. **d)** Group-level cortical projection of model-derived semantic discrepancy feature levels in the control experiment, following the same conventions as in **c**. **e)** Proportion of voxels associated with each feature level within V1, V2, and V3. Bars show the proportion of voxels significantly correlated with low-, mid-, or high-level feature dissimilarity. Voxel inclusion was based on subject index threshold of 0.6. Left panel: main experiment; Right panel: control experiment. In both conditions, low- and mid-level feature representations decrease along the visual hierarchy. However, the proportion of high-level feature-related voxels remains stable in the main experiment but markedly diminishes in the control condition, reflecting reduced reliance on semantic-level expectations. **f)** Subject-level comparison of the proportion of significant vertices in mismatched trials across V1–V3 between the main and control experiments. Each row corresponds to a feature level: low-level (top), mid-level (middle), and high-level (bottom). Boxes represent the proportion of vertices significantly associated with each feature level in individual participants. A significant reduction in the proportion of high-level vertices was observed in V3 under the control condition (p = 0.0391, Wilcoxon signed-rank test, uncorrected). No other comparisons showed significant differences across regions or feature levels.

We then performed voxel-wise linear regression between these feature dissimilarity measures and image-evoked BOLD responses. The resulting maps, projected onto cortical surfaces (Fig. 5c), revealed that: a substantial proportion of vertices in EVC were significantly associated with high-level feature dissimilarity (red), whereas low-level dissimilarity effects (blue) were more concentrated in V1. Vertices sensitive to both types of dissimilarity were highlighted in white. Overall, these spatial distributions indicate that high-level discrepancy-related effects were broadly represented across EVC, whereas low-level effects were more spatially restricted.

To quantify these patterns, we computed the proportion of significant vertices within V1, V2, and V3 for each feature level across 8 participants. We found that in the main experiment, where captions reliably predicted the upcoming images, participants formed explicit high-level semantic expectations. Under a subject index threshold of 0.6, feature dissimilarity analysis revealed that high-level dissimilarity remained relatively stable across early visual cortex: 22.5%, 21.6%, and 20.0% of voxels in V1, V2, and V3, respectively, showing significant correlations with high-level dissimilarity. In contrast, low-level and mid-level dissimilarity effects showed a clear decreasing trend along the visual hierarchy, low-level: 12.8%, 5.3%, 0.2%; mid-level: 27.9%, 14.3%, 12.0% (Fig. 5e).

We then asked whether this pattern depended on the contextual statistics of the task. In the control experiment, where the prevalence of matched and mismatched trials was reversed, high-level dissimilarity-related effects were markedly reduced, particularly in V3 (20.0% to 0.03%), whereas low- and mid-level effects largely retained their hierarchical trends, albeit at somewhat reduced proportions (low-level: 12.2%, 1.5%, 0.0%; mid-level: 17.1%, 14.1%, 7.3%) (Fig. 5d,e).

To further validate this pattern at the individual level, we performed subject-level statistical comparisons on the proportion of significant vertices in the mismatched trials between the two experimental conditions (Fig. 5f). Specifically, there are 400 mismatched trials in the main experiment and 384 mismatched trials in the control experiment. A Wilcoxon signed-rank test revealed a significant reduction in high-level dissimilarity-related vertices in V3 under the control condition (p = 0.0391, Wilcoxon signed-rank test, uncorrected). In contrast, although group-level results in Fig. 5e also indicated reductions in V1 and V2, subject-level differences were not statistically significant (p = 0.313 and p = 0.547, respectively; uncorrected). Consistent with the group-level findings, the proportions of low-level and mid-level dissimilarity-related voxels did not differ significantly across conditions in any of the early visual areas (all p > 0.1, Wilcoxon signed-rank test; uncorrected). Moreover, we demonstrate there is a similar pattern in PPA (Supplementary Fig. 10).

Taken together, these results show that image-evoked responses in early visual cortex are associated with model-derived discrepancy measures at multiple representational levels. In particular, high-level semantic discrepancy-related effects were more prominent in the main experiment and were selectively reduced when contextual statistics were altered, suggesting that caption-defined scene context preferentially shaped high-level response structure in EVC.

### Semantic dissimilarity is associated with reaction time and IFS activity

The visual cortical analyses indicated that caption-defined semantic relationships shaped image-evoked responses. We therefore tested whether these relationships also tracked behavioral performance and activity in higher-order cortex. Using mismatched trials from the main and control experiments, we related trial-wise reaction times (RTs) to model-derived feature dissimilarity and neural activity in higher-order cortical regions.

We first compared RTs between the main and control experiments. Six out of eight participants exhibited significantly slower RTs in the main experiment than in the control (p < 0.001, Mann– Whitney U test, FDR corrected), whereas Subject 1 and Subject 3 showing no significant difference (p = 0.633, Mann–Whitney U test, FDR corrected) (Fig. 6a). All responses occurred within the 3-second image presentation window, confirming that participants responded on the basis of ongoing perceptual processing during image viewing. This pattern aligns with previous literature, which reports faster responses when mismatches are expected^15^ (i.e., in the control experiment, where mismatched trials dominate and are thus expected). In contrast, in the main experiment, where matched trials dominate, mismatched trials constitute a violation of expectation and appear to require more deliberation.

**Figure 6.**
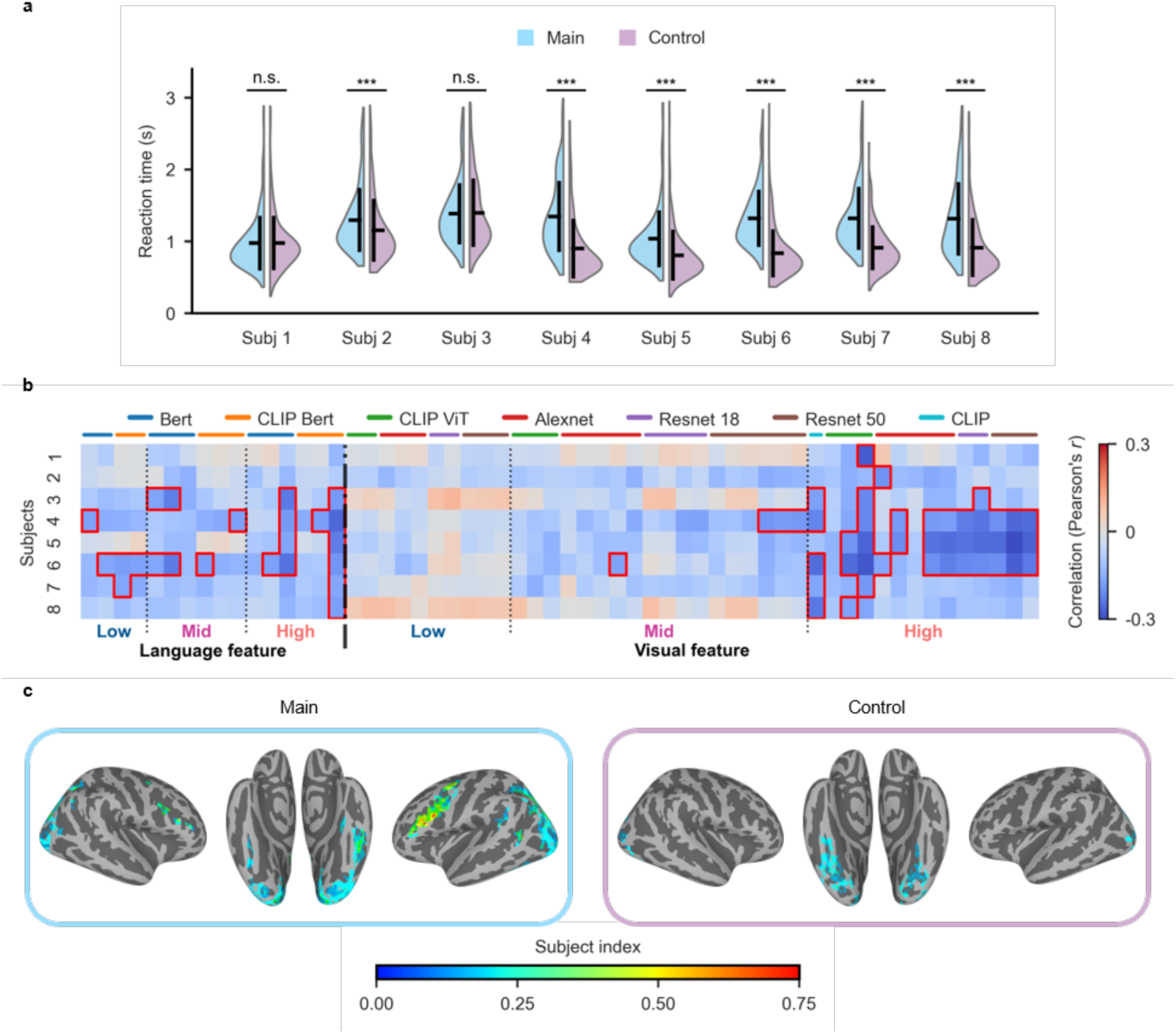
Semantic dissimilarity modulates reaction time and IFS activity. **a)** Reaction time comparisons across eight participants between the main and control experiments. *** indicates p < 0.001 (Mann–Whitney U test, FDR-corrected). **b)** Pearson correlations between reaction time and feature dissimilarity across participants. Horizontal labels indicate the source model of each feature; red boxes highlight significant correlations (p < 0.01, FDR-corrected). Color denotes the Pearson’s r value. Feature levels (low/mid/high) are determined empirically based on model depth, consistent with Fig. 5b. **c)** Subject index (SI) maps showing correlations between BOLD responses and reaction time. Only vertices that showed significant Pearson correlation with reaction time (p < 0.001, uncorrected) at the subject level were projected to the fsaverage cortical surface. Vertices with SI > 0.2 and cluster size > 20 are displayed.

We then asked whether RTs were related to model-derived discrepancy between the caption-defined context and the viewed image. Specifically, we correlated trial-wise RTs with low-, mid-, high-level feature dissimilarity derived from pretrained visual and language models. In the main experiment, all eight participants showed significant negative correlations between RTs and high-level visual dissimilarity (p < 0.01, Pearson correlation, FDR corrected) (Fig. 6b), indicating that larger semantic mismatches were associated with faster responses. Similar trends were observed for high-level language-based dissimilarity, whereas low- and mid-level features showed no consistent relationship with RTs. Importantly, in the control experiment, no significant correlations were observed between RTs and any feature dissimilarity measure, reinforcing the role of semantic expectations in modulating behavior. (Supplementary Fig. 11)

To examine whether this behavioral effect was reflected at the neural level, we next computed voxel-wise correlations between trial-wise RTs and BOLD activity. At the subject level, we identified vertices whose time series correlated significantly with trial-wise RTs (p < 0.001, Pearson correlation, uncorrected). These correlation maps were projected to the average surface space. Notably, we found that IFS showed consistent RT-related activation only in the main experiment and not in the control experiment, even at a relatively lenient threshold (subject index > 0.2, cluster size > 20) (Fig. 6c). These results indicate that the relationship between semantic discrepancy and behavior was accompanied by task-dependent recruitment of IFS.

Taken together, these results reveal a task-dependent relationship among model-derived semantic discrepancy, behavioral performance, and IFS activity. In the main experiment, larger high-level mismatches were associated with faster responses, and RT-related neural activity was consistently observed in IFS. In contrast, these relationships were not evident in the control experiment. This pattern suggests that caption-defined semantic context modulated not only visual cortical responses, but also the coupling between high-level discrepancy structure, behavioral performance, and frontal activity.

### Cross-modal semantic alignment is preferentially represented in suppression regions

We next asked whether the heterogeneous populations identified in early visual cortex differed in their capacity to support cross-modal semantic alignment. To address this question, we developed BrainCLIP (Fig. 7a), a framework inspired by CLIP that maps caption-evoked and image-evoked brain responses into a shared embedding space, allowing us to assess whether neural responses to the same semantic content can be aligned across modalities.

**Figure 7.**
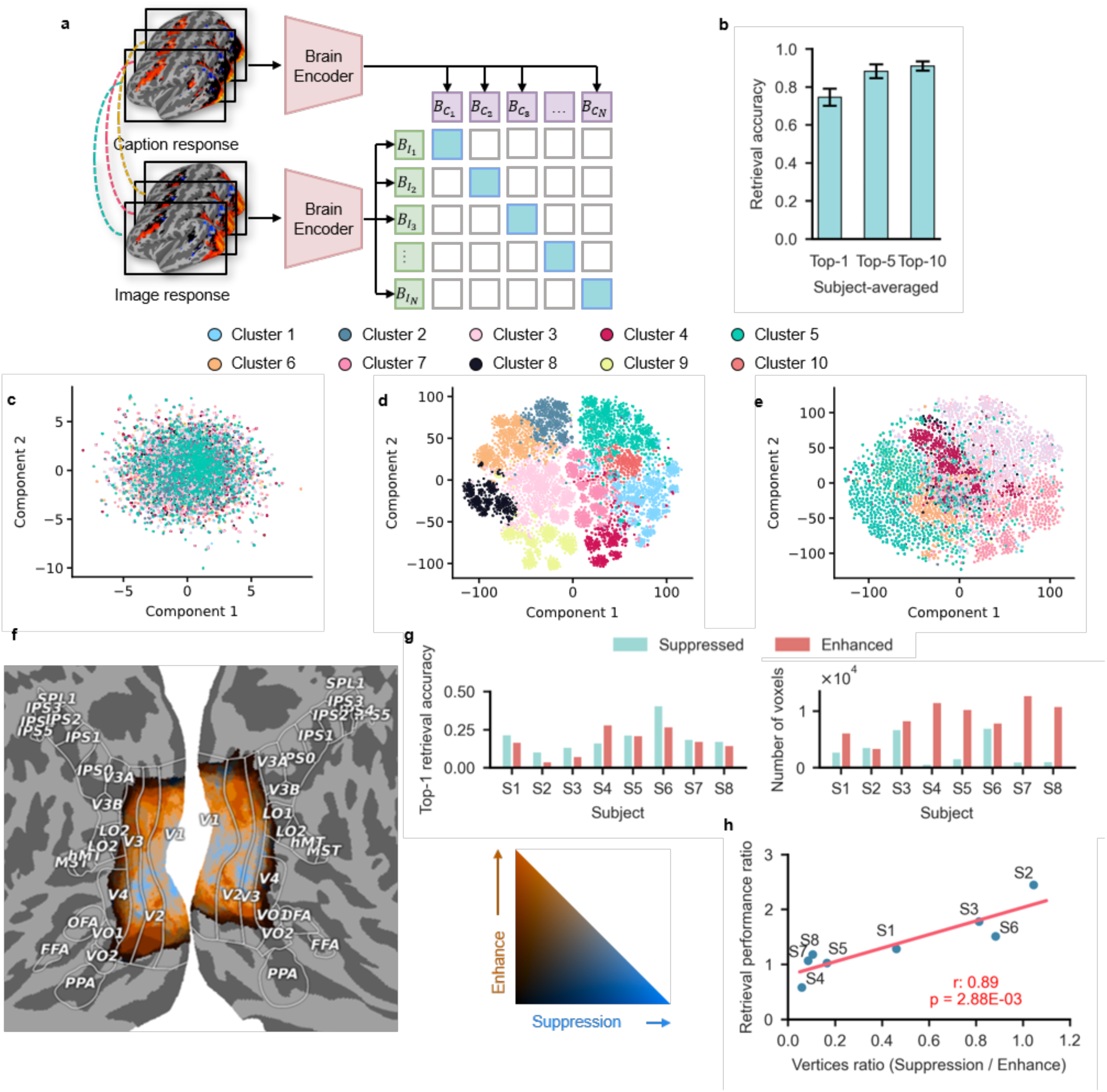
BrainCLIP reveals semantic alignment between brain responses and language-vision embeddings. **a)** Architecture of the BrainCLIP model. Brain responses (beta weights) to semantically matched captions and images were encoded by a shared brain encoder. The model was trained to produce similar embeddings for responses to the same semantic content. **b)** Text retrieval accuracy using BrainCLIP trained on whole-brain responses from 8 participants. Given an image response, the model retrieved the corresponding caption response from a validation set of 400 candidates (chance = 0.0025). The average top-1 accuracy across participants was 74.5%. **c–e**, t-SNE visualizations of brain and model embedding spaces, colored by semantic categories (determined by k-means clustering in CLIP space, k = 10). **c)** t-SNE of raw brain response space for subject 1. **d)** t-SNE of CLIP’s semantic embedding space for subject 1. **e)** t-SNE of BrainCLIP-encoded embedding space for subject 1. **f)** Distribution of suppressed and enhanced voxels in early visual cortex (EVC), projected onto the fsaverage cortical surface. Colors reflect participant-level consistency: blue indicates voxels showing suppression in most participants, yellow indicates voxels showing enhancement. **g)** Performance of BrainCLIP when trained separately on suppression or enhancement regions. Left: text retrieval accuracy across 8 participants; right: corresponding number of voxels used in each region. **h)** Correlation between voxel count and retrieval performance across participants for suppression and enhancement regions. A strong linear relationship was observed (r = 0.89, p = 2.88 × 10^−3^).

We first evaluated BrainCLIP using whole-brain voxel activity. The model successfully performed a cross-modal retrieval task in which image-evoked responses were used to retrieve the corresponding caption-evoked responses of the same semantic content. Across all eight participants, top-1 retrieval accuracy exceeded 70% (mean = 74.55%, standard deviation = 4.51%) (see Supplementary Table 1 for example retrieval results), indicating that brain-wide neural responses contain modality-general semantic structure that can be aligned for cross-modal inference (Fig. 7b). To further evaluate the embedding space, we applied k-means^51^ clustering to CLIP-derived stimulus embeddings, yielding semantic cluster labels, and visualized these using t-SNE^52^. Projecting these cluster labels onto both raw fMRI data and BrainCLIP-transformed embeddings revealed that BrainCLIP more clearly separated stimuli by semantic category, closely mirroring the structure of pretrained CLIP features and demonstrating that our model preserves and enhances semantically meaningful organization in neural responses (Fig. 7c-e).

To understand which brain regions contributed most to successful cross-modal alignment, we leveraged previous results distinguishing suppression and enhancement areas in EVC based on expectation-related signal modulation. These regions were delineated anatomically (Fig. 7f). We then trained separate BrainCLIP models using neural responses from either suppression or enhancement voxels and compared their retrieval performance. Despite using far fewer voxels, models trained on suppression areas achieved retrieval accuracy comparable to, or even exceeding, that of models trained on enhancement areas (Fig. 7g). When retrieval performance was normalized by voxel count as a proxy for information efficiency, suppression voxels consistently showed higher efficiency than enhancement voxels. Moreover, across participants, the relative proportion of suppression voxels was linearly correlated with the relative performance advantage of suppression-based models, suggesting a systematic relationship between predictive structure and representational utility (Fig. 7h).

These results suggest that aligned, cross-modal semantic representations in EVC are not uniformly distributed. Instead, cross-modal semantic alignment was more efficiently expressed in suppression-defined than enhancement-defined regions, consistent with heterogeneous population-level organization of language-guided context effects in EVC. Suppressed signals encode expectation-confirming information and thus provide a compressed and behaviorally useful summary of sensory input. In our task, such predictive structure appears to facilitate semantic alignment across modalities and contributes to improved behavioral performance.

### Population-level representational differences between suppression and enhancement

Building on the finding that suppression regions exhibit stronger cross-modal semantic alignment efficiency than enhancement regions under matched conditions, we next asked how these two populations differ in their representational organization across matched and mismatched trials. To address this, we selected 100 high-response vertices (based on ON-OFF *R*^2^) from suppression and enhancement regions within V2 and V3, while for V1 only suppression regions were included due to insufficient overlap with enhanced regions in some participants (e.g., Subjects 2 and 4). For each region of interest, we constructed voxel-by-trial response matrices (100 voxels × 400 trials) separately for matched and mismatched conditions.

We first asked whether the two populations differed in their ability to discriminate matched from mismatched trials. In suppression-defined regions, multivoxel pattern analysis showed above-chance classification of matched versus mismatched trials, despite lower mean response amplitudes overall (Fig. 8a). A similar effect was also observed in enhancement-defined regions (Fig. 8b), indicating that both populations carried information relevant to semantic condition structure.

**Figure 8.**
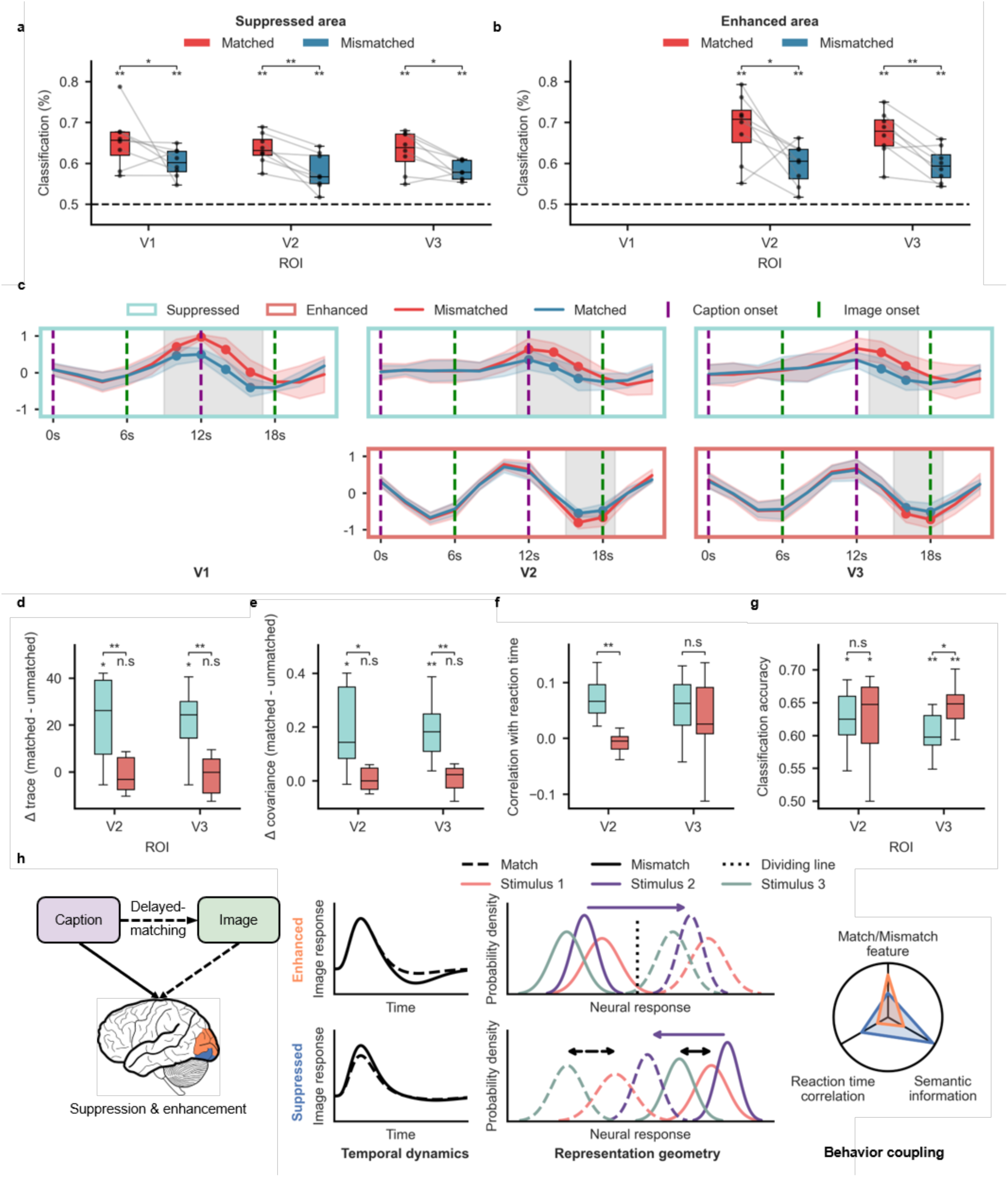
Population-level representational differences between suppression and enhancement regions. **a)** MVPA orientation classification accuracy of semantic matching type in suppression regions. Across V1–V3, classification accuracy was significantly higher for matched than mismatched trials, and exceeded the 50% chance level. **b)** MVPA orientation classification accuracy of semantic matching type in enhancement regions. In V2–V3, classification accuracy was significantly higher for matched than mismatched trials, and exceeded the 50% chance level. **c)** Averaged normalized time courses (mean ± s.d., across 8 participants) within V1–V3, plotted separately for matched and unmatched trials. Time series are scaled to [−1, 1]. Dots indicate time points with significant differences between conditions (p < 0.05, Wilcoxon signed-rank test, FDR corrected). Upper panels: suppression regions; lower panels: enhancement regions. **d)** Group-level comparison of Δ trace values between suppression and enhancement regions within V2 and V3, based on 8 participants. Δ trace was defined as the difference in the trace of the voxel-wise covariance matrix between matched and unmatched conditions. Paired t-tests were used to assess significance. **e)** Group-level comparison of Δ covariance, defined as the difference in mean covariance between matched and unmatched trials, in suppression and enhancement regions. **f)** Correlation between neural responses in suppression and enhancement regions (V2 and V3) and reaction times, showing distinct behavioral coupling across populations. **g)** Classification accuracy of a linear SVM trained to discriminate semantic conditions (matched vs. mismatched) based on multivoxel patterns in suppression and enhancement regions. Results reflect cross-validated decoding accuracy across participants. Chance level was 50%. In enhancement regions, accuracy was not significantly different from suppression region in V2, but significantly exceeded both chance and the suppression region accuracy in V3 (p < 0.05), indicating stronger semantic discriminability in enhanced regions of V3. **h)** Conclusion of population-level difference in 3 different aspects, including temporal dynamics, representation geometry and behavior coupling.

We next examined the normalized time courses within populations. In suppression-defined regions, matched and mismatched responses diverged progressively later along the visual hierarchy, with condition differences emerging at approximately 10 s in V1, 12 s in V2, and 14 s in V3 (image onset at 6 s). Within both V2 and V3, suppression-defined regions showed earlier divergence between matched and mismatched trials than enhancement-defined regions, whose condition differences emerged at around 16 s (Fig. 8c). These results indicate that the two populations differed in the temporal expression of caption-dependent condition effects, with suppression regions responding more rapidly to semantic violations than their enhanced counterparts.

To further characterize population manifold structure, we computed trial-wise covariance matrices for matched and mismatched conditions and compared their traces. We found that suppression regions exhibited significantly greater trace values than enhanced regions (p = 0.008 in V2, p = 0.023 in V3; Wilcoxon signed-rank test, uncorrected) (Fig. 8d), indicating that although mean responses in these areas were lower under matched conditions, their trial-by-trial variance increased. Moreover, the trace values in suppression regions were significantly greater than zero (p = 0.039 in both V2 and V3; Wilcoxon signed-rank test, uncorrected), suggesting a meaningful increase in representational variability under predictive conditions. In contrast, trace values in enhancement regions did not differ significantly from zero in either V2 or V3, indicating more stable population responses despite higher average activity.

In addition, voxel-wise correlations within each neural population exhibited similar patterns: in suppression regions, the correlations among vertices were significantly higher during matched trials compared to unmatched trials (p = 0.046 in V2, p = 0.004 in V3; Wilcoxon signed-rank test, uncorrected) (Fig. 8e), suggesting more consistent information encoding under predictive conditions. In contrast, enhancement regions showed no significant difference in within-region correlations between matched and unmatched trials (p = 0.600 in V2, p = 0.557 in V3), indicating stable response structure regardless of prediction.

We further compared the correlation between RTs and neural activity in suppression versus enhancement regions (Fig. 8f). In V2, suppression regions showed significantly stronger correlations with RT than enhancement regions (p = 0.008, Wilcoxon signed-rank test, uncorrected), suggesting a tighter coupling between neural variability and behavioral performance. No significant difference was observed between the two regions in V3.

To test whether these population-level changes carried behaviorally relevant information, we trained linear support vector machines^53^ (SVMs) to classify matched versus unmatched trials based on the neural response vectors within each ROI (Fig. 8g). Both suppression and enhancement areas showed classification accuracies significantly above chance level (suppression: p = 0.015 in V2, p = 0.002 in V3; enhancement: p = 0.028 in V2, p = 0.002 in V3; Wilcoxon signed-rank test, uncorrected). Moreover, in V3, classification accuracy was significantly higher in the enhanced area compared to the suppression area (p = 0.049, Wilcoxon signed-rank test, uncorrected).

These results demonstrate that both suppression and enhancement regions undergo changes that contribute to multimodal information integration, but along different representational dimensions. Suppression regions enhance semantic discriminability by increasing trial-by-trial variance, thereby supporting finer differentiation of semantic content. In contrast, enhancement regions primarily modulate overall response amplitude, facilitating binary classification of whether inputs match or mismatch predicted semantics (Fig. 8h).

## Discussion

Our findings show that language-defined scene context is incorporated into visual processing before and during image perception, with effects that vary across cortical stage and neural population. Rather than acting only at late interpretive stages, caption-derived semantic context was detectable before image onset, including in early visual cortex, and shaped subsequent image-evoked responses according to the association between the caption and the viewed image. This influence was also related to task performance and frontal engagement, indicating that prior language context was integrated into ongoing task-related processing. Within early visual cortex, contextual effects were not expressed as a uniform modulation. Instead, suppression- and enhancement-defined populations differed in cross-modal alignment, temporal divergence, covariance structure, semantic readout and behavioral coupling. Together, these results suggest that linguistic context enters visual processing as a structured neural state that constrains subsequent scene responses through population-specific interactions, rather than as a single global bias.

An important implication of these findings is that language-defined scene context should not be treated as equivalent to the simpler forms of top-down modulation typically examined in cueing paradigms. Unlike spatial or feature-based cues, a natural-language caption provides abstract, relational, and scene-level semantic constraints that can shape how later visual input is processed without specifying a single low-level sensory template. In this sense, the present paradigm is closer to the problem of how prior knowledge constrains natural scene understanding than to classical cueing designs. At the same time, our results do not isolate a single underlying mechanism. The preparatory and post-stimulus effects observed here may reflect a broader language-driven contextual state that could include contributions from semantic priming, attentional allocation, expectancy, or imagery-like processes. Rather than cleanly separating these components, our results show that language-derived context, whatever its composite mechanism, is expressed within visual cortex in a structured and measurable way before and after image onset.

Our study also helps position language-defined scene context relative to two related lines of work. First, previous studies of visually driven expectation have shown that high-level predictive-error-like signals can be observed even in early visual cortex, indicating that EVC can reflect more than bottom-up sensory input alone^54,55^. Our findings are complementary but conceptually distinct. Rather than using visually specified priors, our task established context through natural language, allowing us to ask how abstract, cross-modal scene meaning shapes subsequent visual processing. In this sense, the present study shifts the focus from visually instantiated expectation to language-defined semantic association. Second, prior work has shown that linguistic context can modulate visual responses, but these effects were largely characterized at the level of overall signal differences^15^. Here, by contrast, we combined model-based analyses of graded caption–image relationships with population-level characterization, allowing us to show not only that language influences visual cortex, but also that this influence tracks semantic structure and is organized across distinct neural populations.

A further contribution of this study is to show that language-related contextual effects in EVC are best understood in terms of complementary population-level organization rather than a single directional modulation. In related work, enhancement and suppression are often examined separately, with emphasis placed on only one side of the effect^56–58^. Our results suggest that this provides only a partial account. Within the same task, suppression- and enhancement-defined populations showed distinct yet complementary functional profiles, differing in cross-modal alignment, temporal divergence, covariance structure, semantic readout, and behavioral coupling. This population-level dissociation is also informative in light of prior encoding-model studies showing that language-aligned models, such as CLIP-like representations, often outperform purely visual models, particularly in higher-level ventral visual regions such as VOTC^16,40^. Our findings are consistent with this broader pattern, but extend it in an important way: rather than locating language-related effects only at the level of regional encoding performance, we show that even within EVC, sensitivity to language-defined scene context is not homogeneous, and is more strongly expressed in specific subpopulations, especially suppression-defined populations. From this perspective, neural population and manifold analyses are valuable not simply because they provide a richer description of the data, but because they reveal how complementary forms of contextual modulation jointly contribute to visual representation.

Despite the strengths of our study, several limitations warrant consideration. First, the temporal resolution of fMRI constrains precise inferences about the dynamics of language-related contextual effects. Although suppressed regions diverged earlier than enhanced regions between matched and unmatched trials, the sluggish hemodynamic response prevents us from determining the precise temporal ordering of these population-level effects relative to ongoing sensory processing. Second, our analyses relied on broad feature spaces derived from pretrained AI models to characterize semantic relationships and cross-modal alignment.

While this approach provided a useful model-based framework for linking language-defined context to neural responses, it does not capture the full fine-grained selectivity of individual voxels or neural populations. Future work combining higher-temporal-resolution recordings with more detailed, voxel-or layer-specific modeling will be important for clarifying both the temporal profile and representational specificity of language-driven contextual influences across cortical hierarchies.

In summary, our study provides a population-level account of how language-defined scene context is represented before and after image onset in human visual cortex. Using a large-scale fMRI dataset and model-based analyses, we show that caption-derived semantic information is detectable prior to visual input, that subsequent image responses are shaped by the semantic relationship between captions and images, and that these effects are organized through distinct neural populations rather than a single uniform modulation. These findings suggest that language does not merely influence late-stage interpretation, but helps structure visual processing itself under natural scene context. More broadly, this work illustrates how large-scale multimodal brain data and AI-inspired analyses can be combined to investigate how abstract knowledge is linked to sensory representation in the human brain.

## Methods

### Subjects

Eight healthy adult participants (4 male, 4 female) took part in this study: S1 (male, age 24), S2 (female, age 27), S3 (male, age 22), S4 (female, age 23), S5 (female, age 23), S6 (male, age 19), S7 (male, age 23), and S8 (female, age 23). All participants were right-handed, had normal or corrected-to-normal vision, and were native Chinese speakers with English as their first foreign language. No special inclusion or exclusion criteria were applied beyond the basic health and eligibility requirements for MRI participation. All scanning sessions were conducted at the AIR Lab, ShanghaiTech University. Prior to participation, all subjects provided written informed consent in accordance with institutional guidelines. The consent form specified the compensation scheme, which included a base rate and an additional bonus awarded upon completion of all scheduled sessions. Participants were informed about the general task structure but were not made aware of the specific proportions of matched versus unmatched trials, nor were they told when repetitions would occur. They were fully informed of the potential duration of the experiment and were reminded of their right to withdraw at any time without penalty. Data quality was assessed continuously during the experiment. Trials or sessions with excessive head motion were excluded from analysis. To mitigate potential effects of long-term memory or repeated exposure, make-up scanning sessions for previously excluded runs were conducted no earlier than three months after the completion of the original set of experiments. All procedures were approved by the Institutional Review Board of ShanghaiTech University.

### MRI data collection

All MRI data were collected at the AIR Lab of the School of Biomedical Engineering, ShanghaiTech University, using a 3T United Imaging uMR 890 scanner equipped with a 64-channel head coil. During scanning, participants wore earplugs that attenuated scanner noise by approximately 30 dB to ensure acoustic comfort. Foam padding was used to stabilize the head and minimize motion artifacts.

T1-weighted anatomical images were acquired using United Imaging’s GRE-FSP sequence with the following parameters: 1 mm³ isotropic voxel size, TR = 8.3 ms, TE = 2.2 ms, inversion time (TI) = 1060 ms, flip angle = 8°, matrix size = 256 × 256, number of slices = 240, field of view (FOV) = 256 mm × 256 mm, and bandwidth = 260 Hz per pixel. The sequence used weak partial echo, no partial Fourier, and was accelerated using United Imaging’s ACS (AI-assisted Compressed Sensing) technology with an acceleration factor of 2. The total acquisition time was 3 minutes and 44 seconds. T2-weighted images were acquired using the United Imaging’s FSE-MX sequence to support cortical surface reconstruction. Parameters were as follows: 1 mm³ isotropic voxel size, TR = 3000 ms, TE = 452.4 ms, matrix size = 256 × 256, number of slices = 240, FOV = 256 mm × 256 mm, and bandwidth = 550 Hz per pixel. The total acquisition time was 3 minutes and 57 seconds.

Functional BOLD images were acquired using a gradient-echo echo-planar imaging (EPI) sequence with the following parameters: 2.5 mm³ isotropic voxel size, TR = 2000 ms, TE = 30 ms, flip angle = 81°, matrix size = 80 × 80, number of slices = 60, FOV = 200 mm × 200 mm, and bandwidth = 2320 Hz per pixel. A multiband factor of 2 was applied to accelerate acquisition, and weak partial Fourier (90%) was used in the phase encoding direction. To correct for susceptibility-induced distortion, functional runs alternated between anterior-to-posterior (AP) and posterior-to-anterior (PA) phase encoding directions, enabling distortion correction during preprocessing.

### Experiment design

The stimuli used in this study were selected from the COCO-CN^34,35,59^ dataset, which provides Chinese-language textual descriptions aligned with natural scene images. To ensure high visual and semantic quality, all images were preprocessed by cropping them into square format using the provided mask annotations, retaining as much of the object content as possible, and then resampled to a standardized resolution of 480 × 480 pixels. Only text– image pairs with high semantic alignment were retained, based on manual inspection. An additional constraint was imposed on the text descriptions: only captions containing 10 to 20 words were included to control for linguistic complexity and reading time. All captions used in the experiment were written in Chinese, and participants were instructed to read silently.

This filtering procedure yielded 10,448 high-quality image-text pairs. From this pool, 9,000 pairs were selected for use in the main experiment. Of these, 1,000 pairs were designated as shared stimuli presented to all eight participants, while the remaining 8,000 were divided such that each participant received 1,000 unique stimulus pairs that were not seen by any other participant.

Each trial consisted of a single text-image pair presented in sequence. Participants first read a text caption for 3 seconds, followed by a 3-second fixation period (central cross), then viewed an image for 3 seconds, followed again by 3 seconds of fixation. Participants were instructed to indicate via button press whether the image matched the preceding caption. A response was only required when the image did not match the text, and this response window spanned the 6–9 second period following trial onset. Throughout the experiment, participants were instructed to maintain fixation on a central cross during all non-stimulus periods and during image presentation, with eye movements minimized.

Each participant viewed 4,000 matching trials in total (1,000 shared + 1,000 unique, each repeated twice), as well as 400 mismatching trials. Unmatched trials were created by randomly pairing text and image stimuli from the full pool of 10,448 items, ensuring minimal semantic overlap. All trials were evenly distributed across 200 fMRI runs. Each run contained 20 matched trials, 2 unmatched trials, and 4 blank trials (6 seconds each), which served as low-level baseline and were randomly interleaved within the run. Blank trials were not allowed to appear consecutively or at the start or end of a run. Each run began with an 8-second fixation and ended with a 4-second fixation period, yielding a total duration of 5 minutes per run. The order of trials within each run was fully randomized.

To validate the semantic specificity of brain responses and to estimate the noise ceiling of decoding models, a separate control experiment was conducted. From the remaining pairs not used in the main experiment, 216 novel image–text pairs were selected. Among these, 184 pairs were assigned as unmatched stimuli and 32 as matched stimuli. For unmatched trials, text–image pairings were generated by randomly shuffling the order of either images or captions from the 184-pair pool, ensuring that no original matches were preserved. Each unmatched combination appeared twice across the control experiment to facilitate noise ceiling estimation. Thus, all participants were exposed to the same 184 images and 184 captions, but the specific pairings in unmatched trials differed due to the randomized mismatch construction. Each participant completed 16 runs, each containing 23 unmatched trials and 2 matched trials. Each run began with an 8-second fixation period and ended with a 4-second fixation, totaling 5 minutes and 12 seconds. In total, each participant viewed 384 mismatched and 32 matched control trials.

Prior to scanning, the display setup was calibrated to ensure that the central fixation cross was located at the center of each participant’s visual field. Visual stimuli were presented at a size subtending approximately 8.2 degrees of visual angle. All stimuli were displayed on a 1920×1080 resolution screen with a 60 Hz refresh rate. Stimulus presentation was controlled using MATLAB and Psychtoolbox^60^, with the average timing error per run verified to be approximately 0.0003 seconds.

Each participant completed at least 30 scanning sessions in total, including 3 sessions for functional localizers and anatomical scans, 25 sessions for the main experiment, and 2 sessions for the control experiment. Each session consisted of 8 runs, resulting in a substantial amount of data per individual.

### Pre-processing of fMRI data

All preprocessing procedures were performed using FSL^61^ (FMRIB Software Library) and FreeSurfer^62^. For each participant, four high-resolution T1-weighted anatomical images were acquired. The brain was first extracted from each T1 image using FSL’s BET tool. These brain-extracted images were then aligned using FLIRT and averaged to enhance the signal-to-noise ratio. The averaged anatomical image was subsequently used as input for FreeSurfer’s recon-all pipeline to reconstruct cortical surfaces. All surface reconstructions were manually inspected and corrected to ensure anatomical accuracy.

Functional MRI data underwent the following preprocessing steps: slice timing correction, distortion correction, motion correction, alignment to anatomical space, one-step resampling, and projection to surface space. Distortion correction was performed using FSL’s topup, based on the mean gradient-echo EPI images from each pair of neighboring runs with opposite phase encoding directions (AP and PA). The estimated susceptibility-induced warp fields were applied to each run individually. For motion correction, each run was registered to the mean image of the first run (after distortion correction) using rigid-body alignment. The reference mean image was then aligned to the anatomical T1 image using boundary-based registration. All transformation matrices and warp fields were concatenated and applied in a single resampling step to minimize interpolation artifacts. The resulting preprocessed functional volumes were finally projected onto each participant’s cortical surface for surface-based analysis.

After preprocessing, we visually inspected the results of distortion correction and evaluated framewise displacement (FD) for motion correction. Runs with excessive motion were excluded from further analysis, and participants were invited back for rescan sessions to ensure sufficient data quality. Once data quality was confirmed, the surface-projected time series were fed into the GLMsingle^63^ toolbox to estimate beta weights for each event. In our model, the caption and the image within each trial were modeled as two separate events to capture their respective neural responses.

To prepare the data for GLMsingle analysis, we followed a normalization strategy described in the original GLMsingle paper for processing the BOLD5000^64^ dataset. Specifically, we first computed the mean signal across all 200 runs. Then, for each session, we estimated a linear transformation (parameters a and b) to rescale the session-level signal so that it matched the across-run average. This linear scaling was applied to reduce inter-session signal variability and improve the stability of GLM estimates. After normalization, we grouped runs containing repeated stimuli from at least two sessions as input to GLMsingle. The model then estimated single-trial beta weights using a regularized general linear model. Finally, within each session, voxel-wise beta weights were z-scored to normalize the signal amplitude and facilitate comparison across sessions and participants.

### Predefined regions of interests

Prior to the main experimental sessions, individual functional localizers were conducted for each of the eight participants to define a set of regions of interest (ROIs). Two independent localizer protocols were used: a population receptive field (pRF) mapping experiment^65,66^ for delineating early visual areas (V1, V2, V3), and a functional localizer (fLoc) experiment^67^ for identifying higher-level category-selective regions (face-, body-, word-, and place-selective areas).

The pRF mapping protocol followed the standard procedures provided by analyzePRF (https://kendrickkay.net/analyzePRF/), using their publicly available stimuli and analysis pipeline. During scanning, participants were instructed to fixate on a central point and respond via button press whenever its color changed. Meanwhile, bar, wedge, and ring stimuli traversed the screen in separate runs. Specifically, each run featured either a multibar or wedgering stimulus configuration, and the two were alternated across four runs (i.e., [multibar, wedgering, wedgering, multibar]) with corresponding AP and PA phase encoding directions ([AP, PA, AP, PA]). Analyses were conducted in cortical surface space, and early visual areas V1, V2, and V3 were manually delineated on each participant’s inflated cortical surface based on polar angle reversals and eccentricity gradients derived from the pRF fits.

To localize high-level visual areas, we used the block-design fLoc localizer developed by the Stanford Vision and Perception Neuroscience Lab (http://vpnl.stanford.edu/fLoc/). Stimuli consisted of images from ten object categories, which were grouped into five semantic domains for ROI definition: faces (adult, child), bodies (body, limb), places (house, corridor), objects (car, instrument), and words (word, number). In each block (duration = 4 s), multiple images from a given category were shown sequentially (500 ms/image), with a 1-back task requiring participants to respond when two identical images appeared consecutively. Each run included 12 blocks per condition, including 12 baseline blocks, all presented in pseudorandom order. Additionally, each run began with a 4-second blank period and ended with an 8-second blank period, allowing for stabilization of the hemodynamic response and capturing post-stimulus activity. Functional data were preprocessed using the minimal preprocessing pipeline described below, and subsequent ROI analysis was performed using AFNI. Face-, body-, place-, and word-selective areas were defined by contrasting each category condition against the rest (e.g., faces > others), thresholded using cluster-level correction at p < 0.05. ROIs were manually labeled on the cortical surface based on the activation clusters in each hemisphere.

### Task-defined ROIs

In addition to functionally localized regions, we defined task-related ROIs based on the neural responsiveness to the main experiment using GLMsingle. Specifically, during the first step of the GLMsingle pipeline, all stimulus events, including both captions and images, were treated as a single condition in an ON-OFF design, to assess voxel-wise responsiveness to the overall task. For each subject, data from at least two sessions were grouped as input to GLMsingle to improve signal stability. This yielded ON-OFF *R*^2^ maps per group, which were then averaged across all groups within each subject. Voxels with averaged *R*^2^ values greater than 1% were defined as task-responsive. These ROIs primarily included large portions of the visual cortex as well as regions in IFS, indicating robust task engagement in both perceptual and higher-order areas.

To further identify regions involved in semantic integration, we examined the differential responses to matched and unmatched trials during image presentation. For each subject, we performed a voxel-wise independent t-test comparing beta weights from matched (n = 4000) and unmatched (n = 400) image trials. Since semantic integration is hypothesized to occur after the image appears, only the image-related beta weights were used. The resulting t-statistic maps were thresholded using cluster-based correction (p < 0.05), and intersected with the task-defined ROIs (based on ON-OFF *R*^2^ > 1%) to obtain individual-level suppression (matched < unmatched) and enhancement (matched > unmatched) regions across the whole brain. To specifically isolate effects within early visual areas, these suppression and enhancement maps were further intersected with early visual cortex (EVC) masks derived from the pRF localizer (union of V1, V2, and V3), yielding EVC-specific suppression and enhancement ROIs.

### Time-course analysis

To characterize the temporal dynamics of semantic suppression effects, we extracted vertex-wise time courses for each subject. For every vertex, single-trial responses were averaged from 0 to 36 seconds (18 TRs) time-locked to the caption onset, separately for matched and unmatched trials. This resulted in a subject-specific matrix of size number of vertices × 2 conditions × 18 time points. Within each subject, the time series were further averaged across vertices within specific ROIs, including V1, V2, V3, and EVC-specific suppression regions.

To quantify the temporal variability of responses, we computed the standard deviation (SD) of the averaged time courses for matched and unmatched conditions within each ROI. We then derived a normalized variability index defined as:

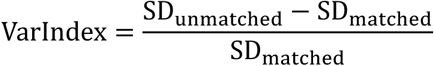

This index was consistently greater than zero across all eight subjects, and the variability in the suppression ROI was significantly higher than that in V1, V2, and V3.

To further characterize functional heterogeneity within the suppression region, we performed unsupervised clustering on the time courses of unmatched trials. For each subject, we applied k-means clustering (k = 3, implemented using sklearn) to the time series of vertices within the suppression ROI. The cluster centroids revealed three dominant temporal profiles: (1) vertices responding selectively to image onset, (2) vertices responding to both caption and image onsets, and (3) vertices showing delayed responses following image onset. We retained the first two clusters for subsequent analysis.

Vertices from the selected two clusters were averaged to yield a representative time course per condition (matched vs. unmatched) per subject. A paired t-test was then performed across subjects at each time point to compare the two conditions. We found that unmatched trials evoked significantly stronger responses than matched trials after image onset, consistent with semantic suppression occurring selectively for expected stimuli.

### Language encoding model

To investigate how semantic information derived from linguistic input is represented in the brain, we developed a voxel-wise language encoding model^40,42,68,69^ based on pre-trained AI language representations. This model was composed of three stages: feature extraction from a transformer-based language model, hierarchical feature mixing, and linear mapping to voxel-level neural responses.

We used the text encoder from Chinese-CLIP^37^, which is based on a RoBERTa-wwm-Large^45^ architecture with 24 transformer layers. For each caption, we extracted both the [CLS] token embedding and the contextual word embeddings from 8 selected layers (layers 1, 2, 4, 8, 12, 16, 20, and 24). Specifically, this yielded a feature set consisting of eight 1024-dimensional [CLS] vectors and eight 51×1024 word embedding matrices (padded to a maximum of 51 tokens to accommodate the typical caption length of 10–20 words).

To reduce computational complexity and facilitate biologically plausible aggregation, we performed a three-step feature mixing procedure. First, all features were passed through a shared 1D convolutional layer (kernel size = 1, depth = 2) to reduce dimensionality from 1024 to 64. Second, for the word embeddings, a 51-dimensional positional attention vector was learned and used to compute a weighted average of token representations within each layer, resulting in eight 64-dimensional word-level summary vectors. Third, a separate attention vector was applied across the eight layers to produce final 64-dimensional representations for both the [CLS] and word-level features. These two were then summed to generate the final mixed linguistic feature vector for each caption.

Importantly, the mixing weights for both the token-level and layer-level aggregation were not shared across the brain but were conditioned on voxel location. Specifically, we learned two lightweight coordinate-to-weight mappings using multi-layer perceptrons (MLPs): one that mapped each voxel’s 3D cortical coordinate to a 51-dimensional token-weight vector, and another that mapped coordinates to an 8-dimensional layer-weight vector. This allowed the encoding model to flexibly learn spatially varying attention patterns over language model representations, reflecting the anatomical specificity of linguistic-semantic processing.

Each voxel was associated with an independent linear regression model that mapped the 64-dimensional mixed feature to its observed fMRI response. To improve generalization and model performance, we incorporated Low-Rank Adaptation^70^ (LoRA) modules into the linear layers of RoBERTa. During training, only the LoRA parameters and downstream feature mixing and regression layers were updated; the original RoBERTa weights were frozen. This setup allowed the model to be fine-tuned in an efficient and data-efficient manner without overfitting the large language model.

To further prevent overfitting and encourage feature diversity, we applied an entropy-based regularization term to the layer mixing weights. This regularizer penalized overly concentrated attention on a single layer, encouraging the model to draw information from multiple levels of representation. The overall loss was defined as a weighted sum of the smooth L1 loss between predicted and actual responses and the entropy regularization term, with the latter weighted by a small constant (1e-6 for subject 1, 2, 3, 5; 1e-5 for subject 6, 7, 8; 1e-4 for subject 4) to balance performance and generalization.

Model training was performed using the AdamW^71^ optimizer with an initial learning rate of 1e-3, a minimum learning rate of 1e-6, and cosine learning rate decay, with the first 2 epochs reserved for learning-rate warm-up. Training proceeded for up to 200 epochs, with early stopping based on validation set performance. If no improvement in voxel-wise *R*^2^ was observed for 20 consecutive epochs, training was terminated.

To further probe which aspects of linguistic input contributed to the model’s predictive power, we performed a series of input manipulations after the encoding model had been fully trained. Since the trained model essentially defines a learned mapping from the space of AI-derived language features to observed neural responses, it can be used to infer how different forms of semantic degradation affect prediction performance. During training, the model was fit using the actual captions viewed by participants along with their corresponding fMRI responses. In the manipulation analysis, we fixed the trained model parameters and evaluated its performance on modified versions of the original captions.

We considered two forms of linguistic perturbation. In the permuted condition, the order of characters within each caption was randomly shuffled, thereby preserving local lexical units but disrupting sentence-level syntactic and semantic coherence. This manipulation allowed us to assess the contribution of compositional meaning while maintaining access to sub-lexical features. In the random condition, each caption was replaced by a syntactically well-formed but semantically unrelated sentence, effectively removing all stimulus-specific information and providing a baseline for evaluating task-relevant encoding. For each of these three input conditions, original, permuted, and random, we computed the voxel-wise *R*^2^ between predicted and observed neural responses, thereby quantifying how encoding performance degraded with the loss of semantic integrity.

### Predictive error modeling

To quantify predictive error signals in the brain during cross-modal semantic processing, we used feature dissimilarity derived from pretrained computer vision models as a proxy for prediction error. A total of 41 distinct features were extracted from pretrained models. We extracted visual features from the vision encoder of Chinese-CLIP (ViT-H/14), AlexNet, and ResNet architectures (ResNet 18 and ResNet 50). Specifically, we used hidden representations from the 1st, 2nd, 4th, 8th, 14th, 20th, 26th, and 32nd layers of ViT-H/14; from all layers (0 through 12) of AlexNet; and from the ReLU activations at each computational block in ResNet18 and ResNet50. In ResNet18, we extracted features from the first two blocks of each residual group, while in ResNet50, features were taken from the first three blocks of each residual group. Following prior literature and architectural conventions, we heuristically grouped all features into three broad representational levels based on their hierarchical position and functional interpretation. Low-level features comprised early ViT layers (1 and 2), the first three layers of AlexNet, and the first residual group in ResNet architectures, which correspond to only one down-sampling operation and primarily encode local contrast and edge-related information. Mid-level features were drawn from ViT layers 4, 8, and 14, mid-stage layers of AlexNet (layers 3 to 7), and the second and third residual groups in ResNet18 and ResNet50, which include multiple down-sampling steps (2 to 3 times) and are thought to represent intermediate visual structures such as shapes and textures. High-level features consisted of the final layers of ViT (layers 20, 26, and 32), the deepest layers of AlexNet (layers 8 to 12), the final residual group in ResNet (which undergoes four down-sampling operations), and the ultimate output of the CLIP visual encoder, all associated with more abstract, object-level or category-level visual representations. For clarity, throughout this study we follow the official PyTorch implementation, where AlexNet is defined as 13 sequential layers (5 convolutional, 5 ReLU, and 3 pooling layers).

For each unmatched trial, participants were presented with a caption followed by an image that violated the semantic content of the caption. To compute a proxy for predictive error, we first identified the image that would have matched the caption (i.e., the semantically congruent image not shown to the participant) and compared it to the actual image shown during the trial. Both the expected (unseen) and actual (seen) images were processed through the pretrained visual models to extract features at each of the 41 selected layers. For each layer, we then computed the cosine distance between the feature vectors of the expected and actual images, using this distance as an operationalization of semantic-level feature dissimilarity, or predictive error, at that level of representation.

To assess the relationship between neural responses and model-derived prediction error, we z-scored both the voxel-wise beta values and the computed feature dissimilarity values. We then performed univariate linear regression at each voxel for each of the 41 feature layers separately. If the regression revealed a statistically significant linear relationship (p < 0.05, uncorrected), the voxel was considered to be significantly modulated by predictive error at that feature level. Finally, each voxel was assigned to one of three representational levels, low-level, mid-level, or high-level, based on which group of features it showed a significant relationship with. Specifically, if a voxel showed significant correlation with at least one low-level feature, it was labeled as sensitive to low-level predictive error; the same logic applied for mid-level and high-level groupings.

### BrainCLIP model

Inspired by the architecture and training principles of CLIP, we developed BrainCLIP to align neural representations of images and text in a shared semantic space. The goal was to examine whether cross-modal brain responses, elicited by either a visual stimulus (image) or a linguistic cue (caption), could be mapped into a common embedding space in which semantically congruent pairs are closer together than incongruent ones.

To achieve this, we first flattened the brain response for each trial (either the image or caption period) into a one-dimensional vector of size *D* (the total number of surface vertices or voxels). Both the image response encoder and the caption response encoder used a shared linear transformation (Linear(*D*, 768)) to project the input vector into a 768-dimensional embedding space. This symmetry ensures that image- and caption-evoked brain activity are mapped using the same representational geometry.

The training objective was to encourage neural responses from semantically matched image-caption pairs to occupy nearby positions in the embedding space, while unmatched pairs were pushed apart. To achieve this, we employed two loss functions: MixCo^72^ and soft-CLIP^73^ loss. MixCo is a contrastive learning strategy that enhances generalization in low-data regimes by mixing training samples and their semantic relationships, and was applied during the first 20 epochs of training. The soft-CLIP loss, which is a differentiable approximation of the standard CLIP contrastive loss, was used throughout all epochs. It computes a temperature-scaled cosine similarity between embeddings and optimizes for alignment of matched pairs while penalizing similarity between unmatched ones.

This training procedure allows BrainCLIP to directly test whether brain activity associated with image and text processing can be linearly aligned in a way that reflects shared semantic content—thus probing the modality-invariance and representational geometry of semantic coding in the human brain.

### Neural population analysis

To examine how predictive modulation influences the structure of population-level neural representations, we conducted a set of analyses within early visual cortical areas. Due to insufficient voxel coverage in specific regions, participant S4 was excluded from all analyses because fewer than 100 suppression-related voxels could be identified in any of the early visual areas. In addition, V1 was excluded from the population-level analysis altogether, as reliable suppression voxels could not be consistently identified across participants. Participant S6, despite limited V1 coverage, met the inclusion criteria in V2 and V3 and was therefore included in the full analysis.

For each of the remaining participants, we selected 100 voxels with the highest task-related *R*^2^ values from the GLMsingle ON-OFF model within both suppression and enhancement regions, separately in V2 and V3. This yielded four voxel groups per participant: suppression-V2, suppression-V3, enhancement-V2, and enhancement-V3. We then extracted response patterns for two types of trials: 400 matched trials that were shared across participants (drawn from shared stimuli and restricted to the first presentation), and 400 unmatched trials from the control experiment. For each participant, this resulted in a data matrix of size 2 (ROIs) × 2 (conditions) × 100 (voxels) × 400 (trials), capturing the trial-by-trial multivoxel activity patterns across both suppression and enhancement regions in V2 and V3.

To replicate previous findings on predictive coding representations, we evaluated the representational geometry associated with semantic prediction within each region of interest (ROI). For each ROI, we analyzed neural responses separately for matched and mismatched conditions, each represented as a 100 voxels × 400 trials matrix. We randomly sampled 320 trials from each condition and concatenated them into a combined dataset. A linear support vector machine (SVM) classifier was then trained to discriminate whether each trial belonged to the matched or mismatched condition, and classification accuracy was computed separately for each condition. This procedure was repeated 30 times with different random subsets to obtain stable estimates of classification performance across participants. Moreover, it is difficult to predict image cluster which is labeled by KNN using CLIP features (Supplementary Fig. 12 & 13).

We analyzed this data using three complementary measures to assess differences in neural population coding between conditions. First, we computed the element-wise difference between the voxel-by-voxel covariance matrices of matched and unmatched trials and calculated its mean value, which reflects the overall change in coactivation structure. Second, we quantified the trace of the covariance difference matrix, which summarizes changes in population-wide variance and inter-voxel dependency. Third, to assess the separability of multivoxel patterns between matched and unmatched conditions, we combined all 800 trials and trained a linear SVM to classify trial types based on population responses, using five-fold cross-validation to evaluate classification accuracy. Together, these analyses allowed us to characterize how semantic expectation modulates not only the strength but also the geometry and informational content of distributed neural representations in early visual cortex.

### Reporting summary

Further information on research design is available in the Nature Research Reporting Summary linked to this article.

## Acknowledgments

This work is supported by the National Science and Technology Major Project of China (2025ZD0217000, Y.L.), National Natural Science Foundation of China (32371154, Y.L.), Shanghai Rising-Star Program (24QA2705500, Y.L.), the Lin Gang Laboratory (LG-GG-202402-06 and LGL-1987-18, Y.L.), and the State Key Laboratory of Advanced Medical Materials and Devices (YGSKL2026ZZ01 to Y. L.). The computations in this research are supported by the HPC Platform of ShanghaiTech University.

## Author contributions

Conceptualization: Y.L., S.G., and R.-Y. Z.; Methodology: S.L., S.G., R.-Y.Z. and Y.L.; Software: S.L.; Investigation: S.L., Z.J., R.-Y.Z., and Y.L.; Data Curation: S.L., Z.J. and R.-Y.Z.; Formal analysis: S.L. and Y.L.; Resources: Y.L.; Writing – Original Draft: S.L.; Writing – Review & Editing: S.L., S.G., R.-Y.Z. and Y.L.; Supervision: Y.L.

## Data availability

Raw fMRI Data and the derivatives are available at the Science Data Bank (https://doi.org/10.57760/sciencedb.27580). The data to reproduce the figures in this study are available at https://osf.io/gj3qx/.

## Code availability

Custom code is available at https://github.com/lishurui0612/cross_modal_integration.

