## Supplementary Information for "Linguistic Context Shapes Representations in the Human Visual Cortex"

### Table of Contents

**Supplementary Figure 1.** Two example voxels illustrating GLM fitting performance.

**Supplementary Figure 2.** Similarity matrices across trials for different beta estimates produced by GLMsingle.

**Supplementary Figure 3.** Relationship between caption- and image-evoked responses in randomly selected voxels.

**Supplementary Figure 4.** Distribution of caption–image correlation and variance inflation factor across voxels.

**Supplementary Figure 5.** Eye movement patterns during caption and image viewing in matched and mismatched trials.

**Supplementary Figure 6.** Screen-space pRF distribution of the EVC suppression region.

**Supplementary Figure 7.** Group-level distribution of enhancement voxels across the whole brain.

**Supplementary Figure 8.** Averaged normalized time courses (mean ± s.d., across 8 participants) within two ROIs (FFA & PPA), plotted for matched and unmatched trials.

**Supplementary Figure 9.** Prediction performance ($R^2$) across the three caption conditions for each of the 8 participants using RoBerta.

**Supplementary Figure 10.** Subject-level comparison of the proportion of significant vertices across FFA & PPA between the main and control experiments.

**Supplementary Figure 11.** Pearson correlations between reaction time and feature dissimilarity across participants in control experiment.

**Supplementary Figure 12.** Five-way image classification across eight participants. Group-averaged decoding accuracy for five image clusters.

**Supplementary Figure 13.** Five-way image classification of eight participants.

**Supplementary Table 1.** Example results of retrieval performance for subject 1.

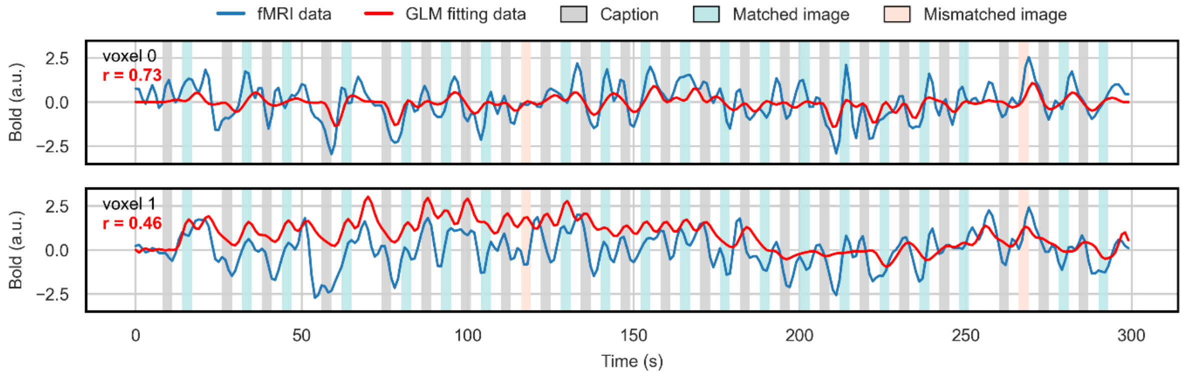


**Supplementary Figure 1. Two example voxels illustrating GLM fitting performance.** The blue curve represents the minimally preprocessed fMRI data, and the red curve represents the GLM-predicted signal. Shaded blocks indicate stimulus duration: gray for captions, blue for matched images, and red for mismatched images.

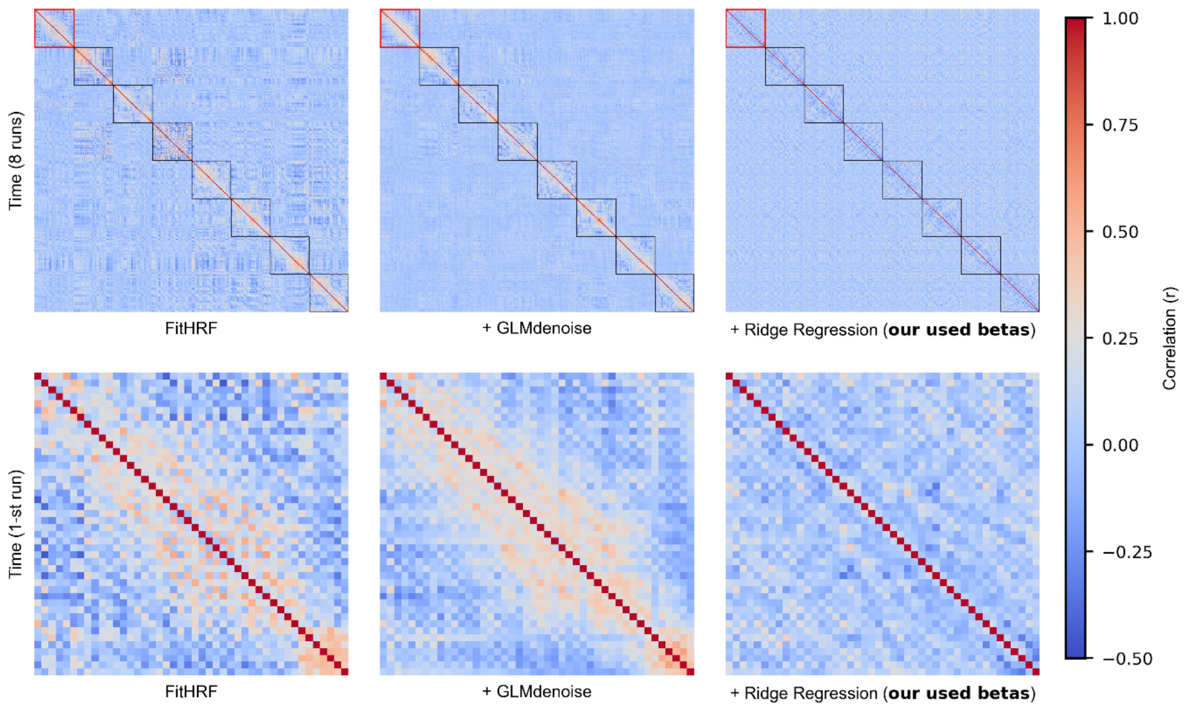


**Supplementary Figure 2. Similarity matrices across trials for different beta estimates produced by GLMsingle.** The first row shows the trial-by-trial similarity matrices for all eight runs from Subject 3, Session 1; each black box denotes one run. The second row shows an enlarged view of the first run. In the beta version used in the main analyses, the similarity between adjacent caption- and image-evoked responses is much lower, supporting the temporal separation of the two signals.

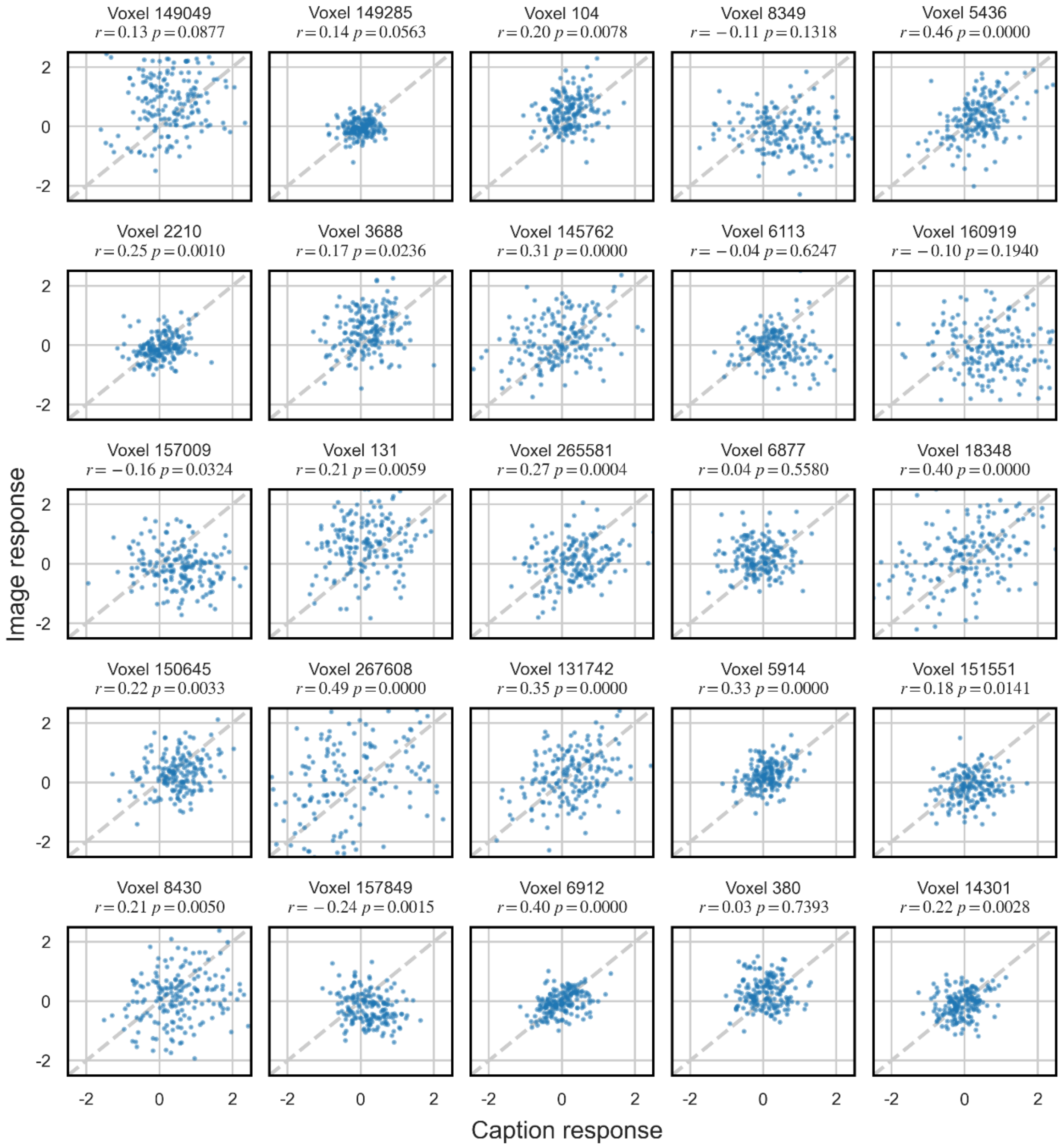


**Supplementary Figure 3. Relationship between caption- and image-evoked responses in randomly selected voxels.** For 25 randomly selected voxels, scatter plots were generated for a single trial, with the caption response plotted on the x-axis and the image response plotted on the y-axis, and the correlation between the two responses was computed. There was little evidence of negative correlation between caption- and image-evoked responses, arguing against the possibility that temporally adjacent activations led to a sustained increase in the BOLD signal, thereby causing overestimation of the caption response and compensatory underestimation of the image response. In addition, most correlations ranged from 0.1 to 0.3, suggesting that collinearity between the two variables was not severe.

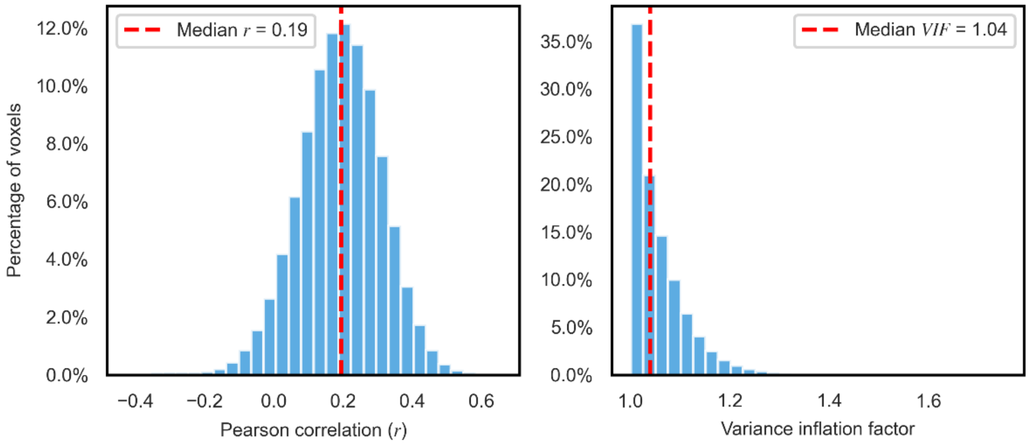


**Supplementary Figure 4. Distribution of caption–image correlation and variance inflation factor across voxels.** Histograms of the correlation between caption- and image-evoked responses across voxels, together with the distribution of variance inflation factor (VIF) values. The mean caption–image correlation was r = 0.19, and the mean VIF was 1.04, indicating low collinearity between the two regressors.

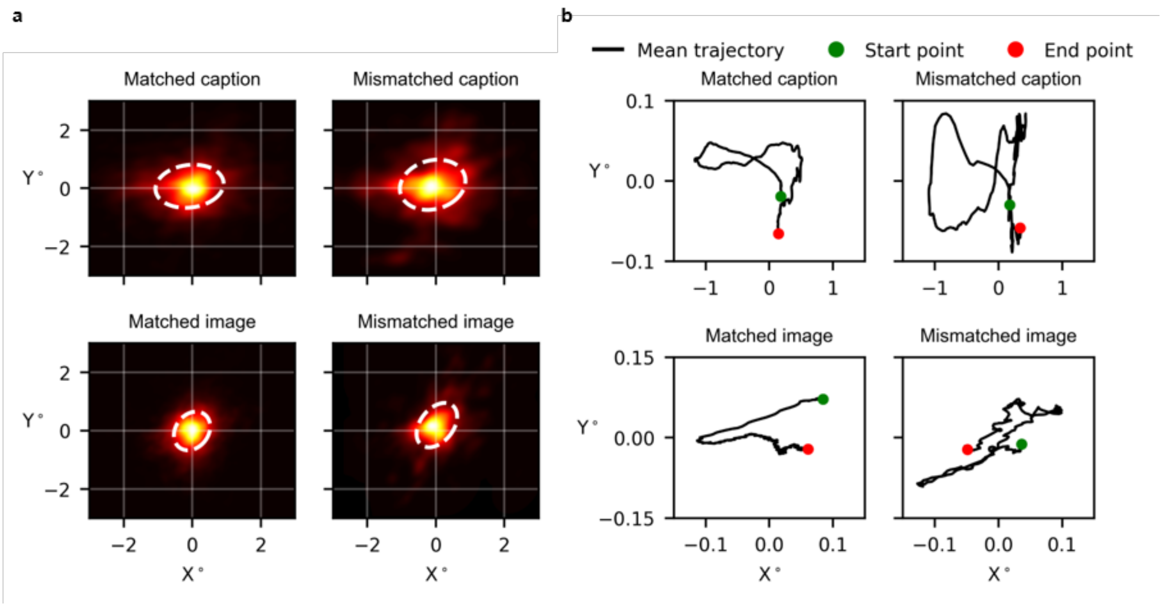


**Supplementary Figure 5. Eye movement patterns during caption and image viewing in matched and mismatched trials. a)** Eye-tracking heatmaps for four conditions: captions and images in matched trials, and captions and images in mismatched trials. White dashed lines indicate the mean ± 1 SD of gaze position. During caption presentation, a clear horizontal eye-movement pattern consistent with reading was observed. During image presentation, gaze remained centered in both matched and mismatched trials. **b)** Mean eye-movement trajectories under the same four conditions. During caption presentation, variation along the y-axis was minimal, whereas pronounced horizontal shifts along the x-axis reflected text reading. During image presentation, gaze remained largely confined within ±0.1° around the fixation center in both conditions. Notably, the image display extended across a much larger range of ±4°.

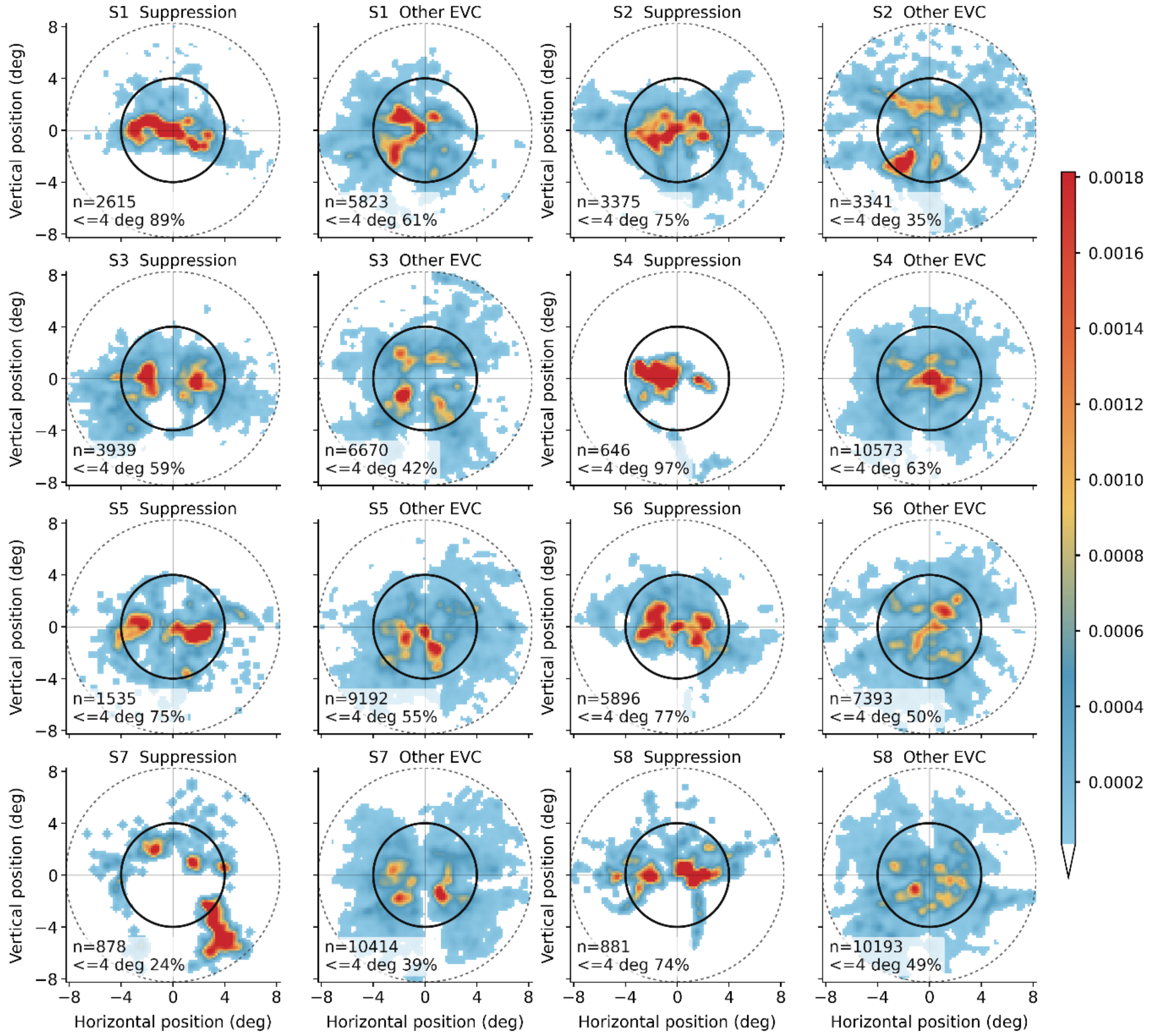


**Supplementary Figure 6. Screen-space pRF distribution of the EVC suppression region.** For each participant, screen-space pRF density maps are shown separately for EVC suppression vertices and the remaining EVC vertices. Only vertices with reliable retinotopic fits were included ($R^2 > 3$). Color indicates the normalized pRF density within each ROI. The solid circle marks the 4° stimulus display range, and the dashed circle marks the retinotopy mapping aperture. Text in each panel reports the number of reliable vertices and the fraction falling within the 4° display range. The suppression region is biased toward central visual space, although it is not exclusively confined to the foveal display range.

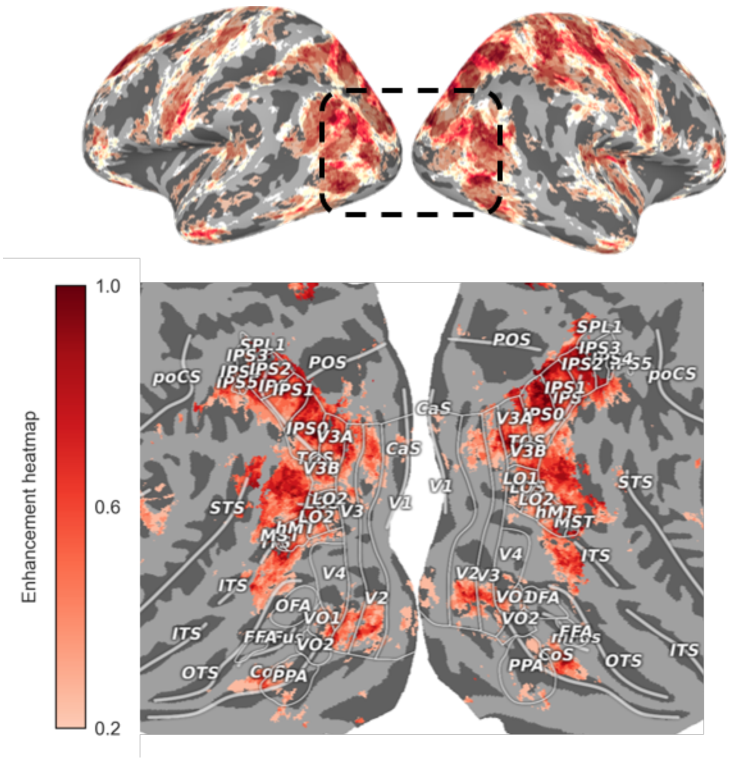


**Supplementary Figure 7. Group-level distribution of enhancement voxels across the whole brain.** The boxed region highlights the dorsal visual cortex, with a magnified view shown below. The enhancement heatmap reflects the number of participants exhibiting enhancement at each voxel after alignment to the standard cortical template.

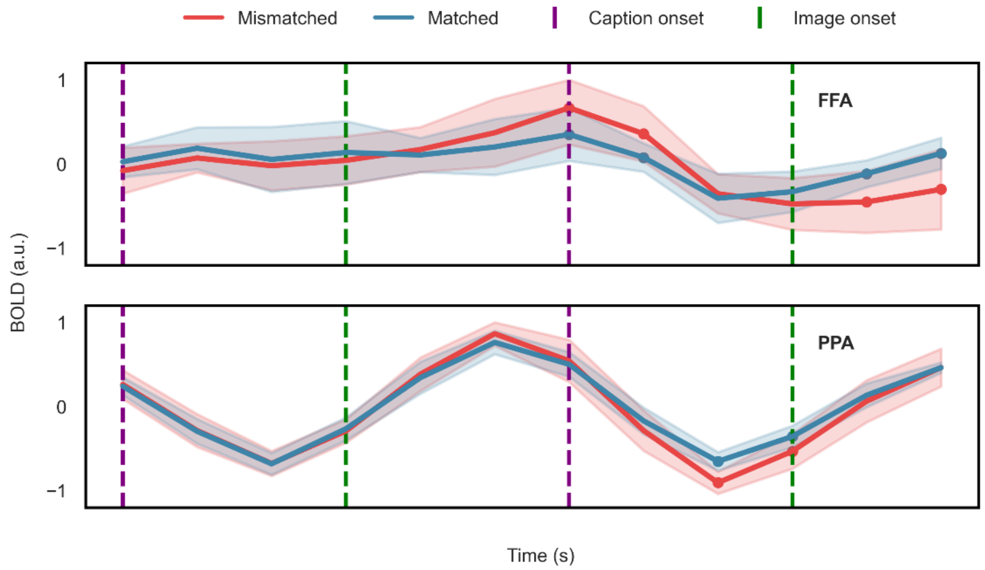


**Supplementary Figure 8. Averaged normalized time courses (mean ± s.d., across 8 participants) within two ROIs (FFA & PPA), plotted for matched and unmatched trials.** Time series are scaled to [−1, 1]. Dots indicate time points with significant differences between conditions ($p < 0.05$, Wilcoxon signed-rank test, FDR corrected).

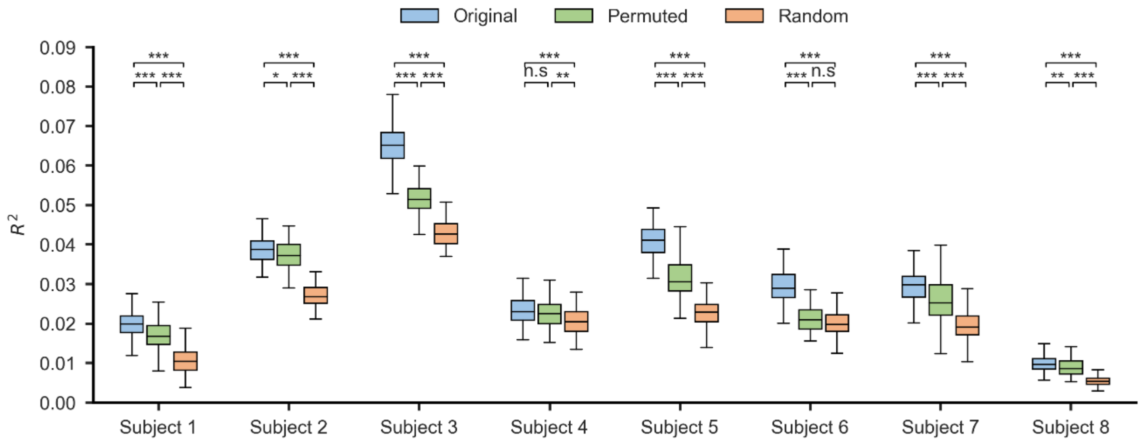


**Supplementary Figure 9. Prediction performance ( $R^2$ ) across the three caption conditions for each of the 8 participants using RoBerta.** Asterisks indicate statistical significance across conditions (n.s.: not significant, *: p < 0.05, **: p < 0.01, ***: p < 0.001; Mann–Whitney U test, Bonferroni correction).

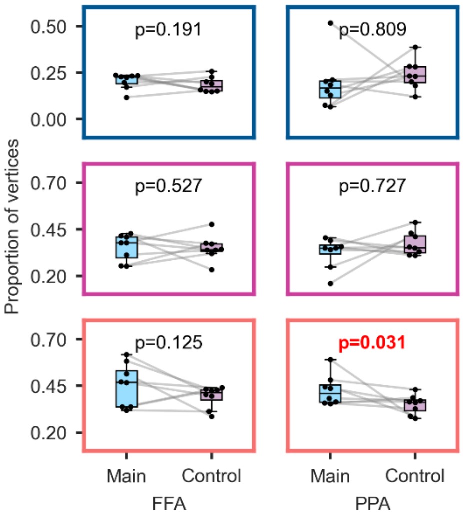


**Supplementary Figure 10. Subject-level comparison of the proportion of significant vertices across FFA & PPA between the main and control experiments.** Subject-level comparison of the proportion of significant vertices across FFA & PPA between the main and control experiments. Each row corresponds to a feature level: low-level (top), mid-level (middle), and high-level (bottom). Boxes represent the proportion of vertices significantly associated with each feature level in individual participants. A significant reduction in the proportion of high-level vertices was observed in PPA under the control condition ($p = 0.031$, Wilcoxon signed-rank test, uncorrected). No other comparisons showed significant differences across regions or feature levels.

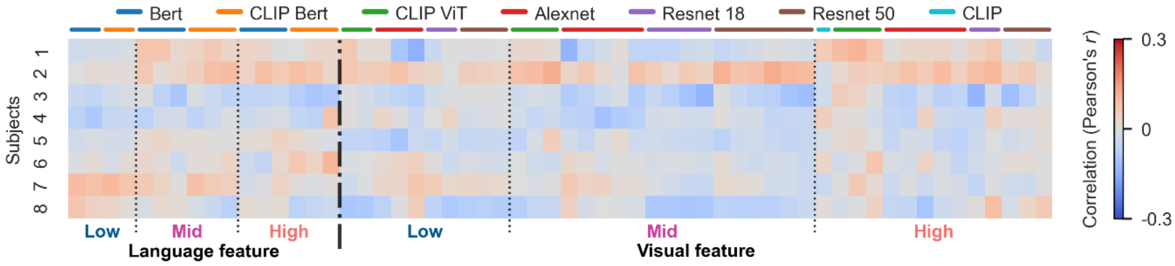


**Supplementary Figure 11. Pearson correlations between reaction time and feature dissimilarity across participants in control experiment.** Horizontal labels indicate the source model of each feature; red boxes highlight significant correlations ($p < 0.01$, FDR-corrected). Color denotes the Pearson's r value. Feature levels (low/mid/high) are determined empirically based on model depth, consistent with Fig. 4b.

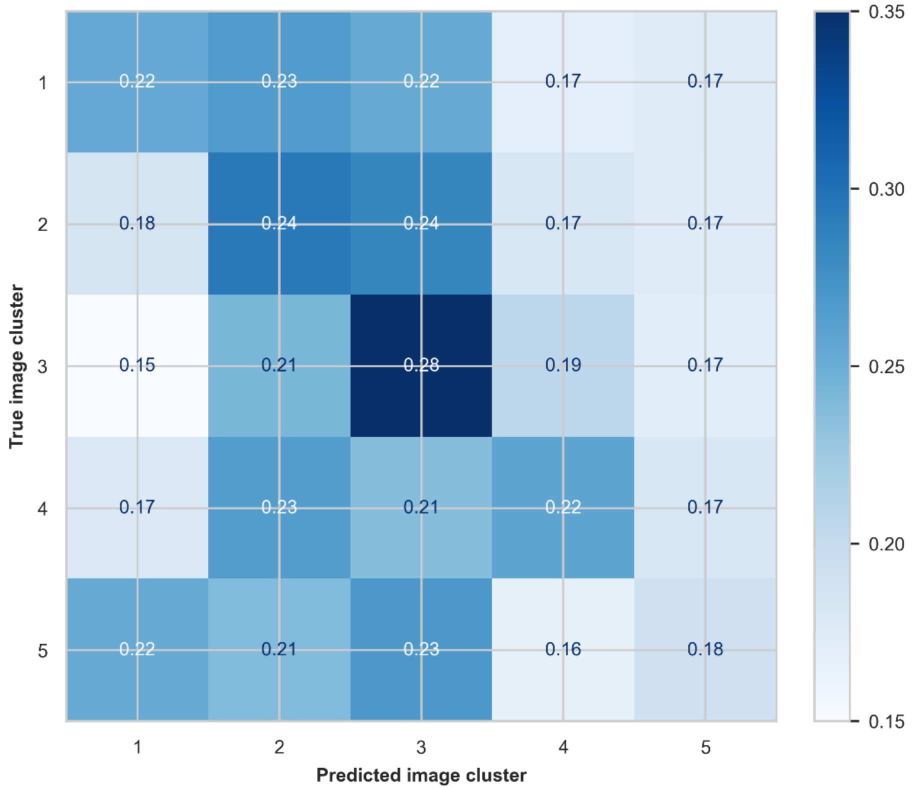


**Supplementary Figure 12. Five-way image classification across eight participants. Group-averaged decoding accuracy for five image clusters.** The results indicate that neural responses in this task could not reliably distinguish between image categories, suggesting minimal category-selective information under the current paradigm.

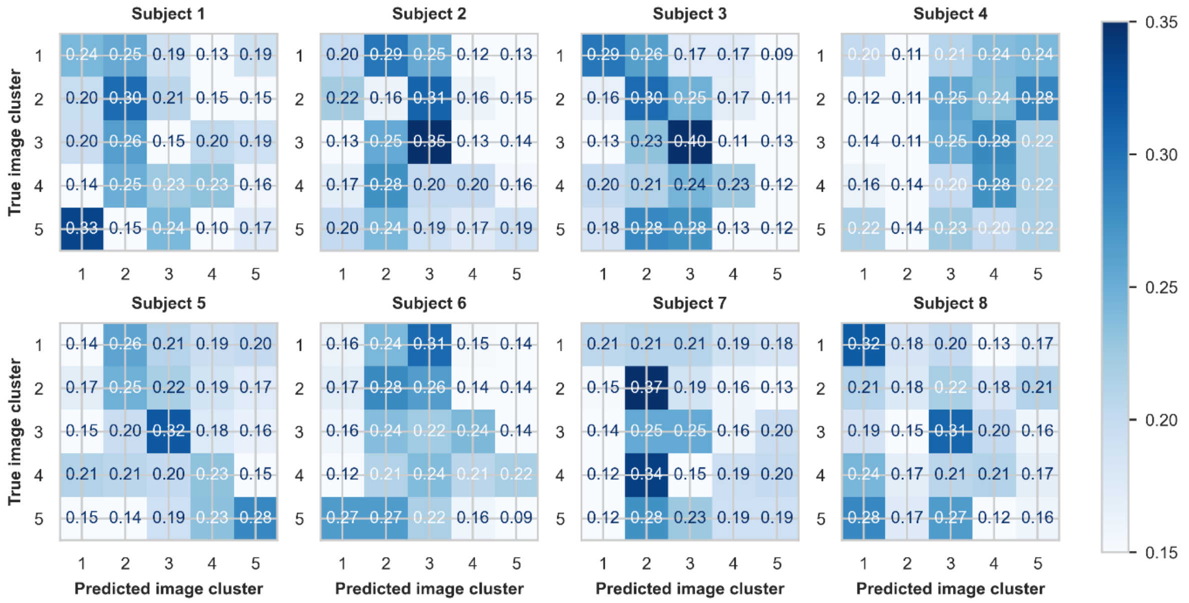


**Supplementary Figure 13. Five-way image classification of eight participants.** The results indicate that neural responses in this task could not reliably distinguish between image categories, suggesting minimal category-selective information under the current paradigm.

| Image stimulus | Corresponding caption | Retrieval top-1 | Retrieval top-2 | Retrieval top-3 |
|---|---|---|---|---|
| 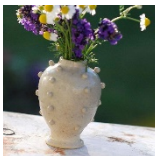 | A vase filled with beautiful little flowers | A vase filled with beautiful little flowers | A boy is playing skateboarding, jumping in the air | A group of zebras stood on the grassland |
| 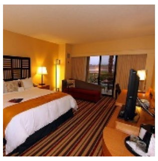 | A clean and warm room | A white plush bear doll is placed in front of the laptop | A clean and warm room | A clean and warm room |
| 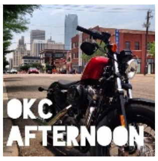  | A brand-new motorcycle parked by the roadside | A brand-new motorcycle parked by the roadside | A brand-new motorcycle parked by the roadside | A black cow stands by the wire fence |
| 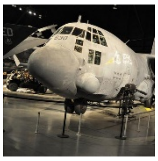 | An old-fashioned airplane is on display in the museum | An old-fashioned airplane is on display in the museum | A large bus is parked on the roadside | An old-fashioned airplane is on display in the museum |
| 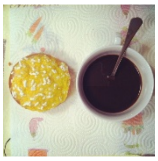 | There is a plate of cake and a cup of coffee on the table | There are many different types of donuts in the box | Two giraffes playing under the stone | A wooden cart was piled with green bananas |

**Supplementary Table 1. Example results of retrieval performance for subject 1.** This table presents five example retrieval cases based on the whole-brain BrainCLIP model. For each visual stimulus (image), we show the top-3 retrieved captions ranked by cosine similarity. Ground-truth captions (i.e., the true paired captions from the original dataset) are also listed for reference. Example images are derived from the COCO dataset (Lin et al., 2014).
